# Epigenetic modulation in the pathogenesis and treatment of inherited aortic aneurysm conditions

**DOI:** 10.1101/2021.02.12.431010

**Authors:** Benjamin E. Kang, Rustam Bagirzadeh, Djahida Bedja, Jefferson J. Doyle, Elena G. MacFarlane, Harry C. Dietz

## Abstract

Shprintzen-Goldberg syndrome (SGS) is a rare systemic connective tissue disorder characterized by craniofacial, skeletal, neurodevelopmental, cutaneous, and cardiovascular manifestations, including aortic root aneurysm. It has significant phenotypic overlap with both Marfan syndrome (MFS) and Loeys-Dietz syndrome (LDS). We previously reported that SGS is caused by heterozygous mutations in the Sloan-Kettering Institute proto-oncogene (*SKI*), which encodes a potent suppressor of transforming growth factor beta (TGFβ) target gene expression. Herein, we show that mouse lines harboring orthologous amino acid substitutions in *Ski* recapitulate multiple human SGS phenotypic manifestations, including skin collagen deposition, skeletal kyphosis, behavioral hypoactivity, and aortic root aneurysm. Furthermore, aortic root aneurysm in SGS mice is associated with both increased acetylation of histone H3 at lysine-27 (H3K27) and TGFβ target gene expression, all of which can be ameliorated by pharmacological CBP/P300 inhibition in vivo; similar findings were seen in cultured dermal fibroblast from SGS patients. Aortic root growth is also abrogated in a mouse model of MFS by selective CBP/P300 inhibition in association with blunted expression of TGFβ target genes. These data document excessive H3K27 acetylation and hence TGFβ target gene expression in the pathogenesis of inherited presentations of aortic root aneurysm and the therapeutic potential of pharmacological epigenetic modulation.

## Introduction

Shprintzen-Goldberg syndrome (SGS) is a rare, autosomal dominant systemic connective tissue disorder (CTD) characterized by craniosynostosis, severe skeletal deformities, aortic root dilatation, minimal subcutaneous fat, intellectual disability, and neurodevelopmental anomalies (1, 2). It shows considerable phenotypic overlap in the craniofacial, skeletal, skin and cardiovascular systems with both Marfan syndrome (MFS) and Loeys-Dietz syndrome (LDS), with the additional findings of mental retardation and severe skeletal muscle hypotonia.

Mutations in the *FBN1* gene encoding the extracellular matrix protein fibrillin-1 cause MFS, while heterozygous mutations in the genes encoding TGFβ ligands (*TGFB2* or *TGFB3*), receptor subunits (*TGFBR1*or *TGFBR2*) or intracellular signaling intermediates (*SMAD2* or *SMAD3*) cause LDS (3, 4). This has led to the recognition that dysregulated TGFβ signaling plays a role in the pathogenesis of both MFS and LDS (4, 5). In prior work, we have demonstrated increased activation of both canonical (i.e. Smad2/3) and non-Smad (i.e. MAPK; Erk1/2) TGFβ-dependent signaling cascades in affected tissues of MFS and LDS mouse models (5-8). Furthermore, multiple phenotypic manifestations in mice, including aortic root aneurysm, can be ameliorated by postnatal administration of TGFβ-neutralizing antibody, and/or the angiotensin II (Ang-II) type 1 (AT1) receptor blocker (ARB) losartan, in association with blunted Smad2/3 and Erk1/2 activation (7, 9-11). We also demonstrated that genetic ablation of *Smad2* in cardiac neural crest-derived vascular smooth muscle cells could abrogate aortic root aneurysm formation in mice with LDS caused by a kinase domain mutation in *Tgfbr1* (12). This occurred in association with normalization of the tissue signature for excessive TGFβ signaling in the aortic wall.

Despite these observations, there were a number of controversies that derived from this initial work. First, certain features of SGS, including craniosynostosis, altered palatogenesis, and aortic aneurysm, have variably been associated with low TGFβ signaling states (13-15). This raised the question as to whether low or high TGFβ signaling drives disease pathogenesis in SGS and related CTDs. This has been further brought into question by the observation that LDS is predominantly caused by heterozygous missense substitutions affecting the kinase domain of either TGFβR1 or TGFβR2, that can cause a context-specific decline in TGFβ signal propagation in cell culture systems (4, 16). Furthermore, LDS-like phenotypes are caused by heterozygous loss-of-function mutations in *SMAD3,TGFB2*, or *TGFB3*, all positive effectors of TGFβ signaling (17, 18). Once again, haploinsufficiency for these genes associated with a tissue signature for high TGFβ signaling including excessive phosphorylation and nuclear translocation of Smad2/3 and enhanced expression of typical TGFβ target genes including *COL1A1, COL3A1, CTGF, MMP2* and *MMP9* (17, 19, 20). Taken together, these seemingly contradictory data have engendered considerable controversy regarding the precise role of TGFβ signaling in the pathogenesis of inherited forms of thoracic aortic aneurysm.

Given the extensive phenotypic overlap of SGS with both MFS and LDS, we hypothesized that aberrant TGFβ activation likely underlies SGS and that identification of the genetic basis of the syndrome would likely inform our understanding of both it and related CTDs. We and others previously identified that de novo heterozygous mutations in the receptor-activated SMAD (R-SMAD) binding domain of the Sloan-Kettering Institute proto-oncogene (*SKI*), a known repressor of TGFβ signaling, cause SGS (21, 22). Furthermore, primary dermal fibroblasts from SGS patients grown at steady-state showed a cell-autonomous increase in transcriptional output of many TGFβ–responsive genes. These data supported the conclusion that SGS pathogenesis appears to be driven by high TGFβ signaling, presumably from loss of suppression by mutant SKI protein. It remains unclear, however, whether these mutations in *SKI* drive SGS pathogenesis via a loss-of-function, dominant-negative, or gain-of-function mechanism.

Because of the plethora of SKI binding partners, the downstream consequences of *SKI* mutations remain uncertain. SKI appears to bind to an array of partners, including R-SMADs (SMAD2 and SMAD3), SMAD4, SKI itself (during dimerization), as well as SKI-like peptide (SKIL), and transcription factors such as CBP/P300, mSin3A, SNW1, N-CoR and HDAC1 (23-30). SGS-causing mutations have to date clustered in the R-SMAD binding domain of SKI towards the N-terminal end of the protein, including substitution of an amino acid that has previously been shown to be essential for SKI-SMAD3 interactions (e.g. p.Leu21Arg) (30). Interestingly, a version of SKI lacking this R-SMAD binding domain retained its ability to regulate SMAD-mediated transcriptional activation in a transient transfection-based reporter system, but failed to dissociate CBP/P300 from the SMAD complex (28, 30), suggesting that CBP/P300 could play a role in SGS. More recently, it was shown that regulated degradation of SKI requires interaction with SMAD2 or SMAD3 and SMAD4; SGS mutations that prevent SMAD2/3 binding resulted in increased stability and hence abundance of mutant SKI, which retained the ability for transcriptional repression of some TGFβ target genes, as evidenced by reduced induction of these transcripts in cultured cells expressing mutant SKI to acute stimulation with exogenous TGFβ (31). Many questions remain regarding the potentially opposing influences of different aspects of altered SKI homeostasis in SGS including increased stability and abundance that is perhaps offset by altered efficiency for recruitment to regulatory elements of target genes. Given that SGS-causing mutations in SKI protein interfere with binding to pSMAD2 and pSMAD3, altered transcriptional regulation is presumably mediated through interaction with SMAD4 only and by aberrant regulation of epigenetic modulators such as CBP/P300. The net effect of heterozygous SGS mutations on TGFβ signaling may vary based on cell type, baseline signaling status, redundancy of autoregulatory factors, and the potential for chronic compensatory events.

In an attempt to inform these questions, we generated and characterized a knock-in mouse model of SGS and explored the molecular mechanisms driving disease pathogenesis in these animals. We used this information to develop a novel therapeutic strategy for the disorder, which also shows efficacy in a well-characterized mouse model of MFS. Finally, we show concordant molecular events and therapeutic potential in cultured dermal fibroblasts from patients with SGS.

## Results

### SGS mouse models recapitulate the phenotype of patients with SGS

To determine whether mutations in SKI are sufficient to recapitulate the SGS phenotype in mice, several targeted mouse lines were developed (Fig. 1A). *Ski*^+/−^ mice are heterozygous for a deletion of exons 2 and 3 of *Ski*, which leads to functional haploinsufficiency due to nonsense-mediated mRNA decay. *Ski*^G34D/+:Neo^ mice are heterozygous for a missense mutation (p.Gly34Asp) previously observed in a patient with severe SGS (21); this allele retains the neomycin resistance cassette, which causes transcriptional interference and also leads to functional haploinsufficiency. *Ski*^G34D/+^ mice are generated by breeding of *Ski*^G34D/+:Neo^ mice to transgenic mice expressing an ubiquitous Cre recombinase (CMV-Cre), which eliminates the Neo cassette, thus allowing transcription of the mutant allele and expression of mutant SKI. As expected, both haploinsufficient lines expressed half the normal complement of *Ski* mRNA, while the heterozygous *Ski*^G34D/+^ line expressed significantly higher levels of *Ski* mRNA, to levels indistinguishable from control mice (Fig. 1B). The *Ski*^G34D/+^ mouse line uniquely expressed the mutant allele, as shown by a diagnostic BamH1 restriction fragment of amplified cDNA that was present only in this line (Fig. 1C).

**Figure 1.**
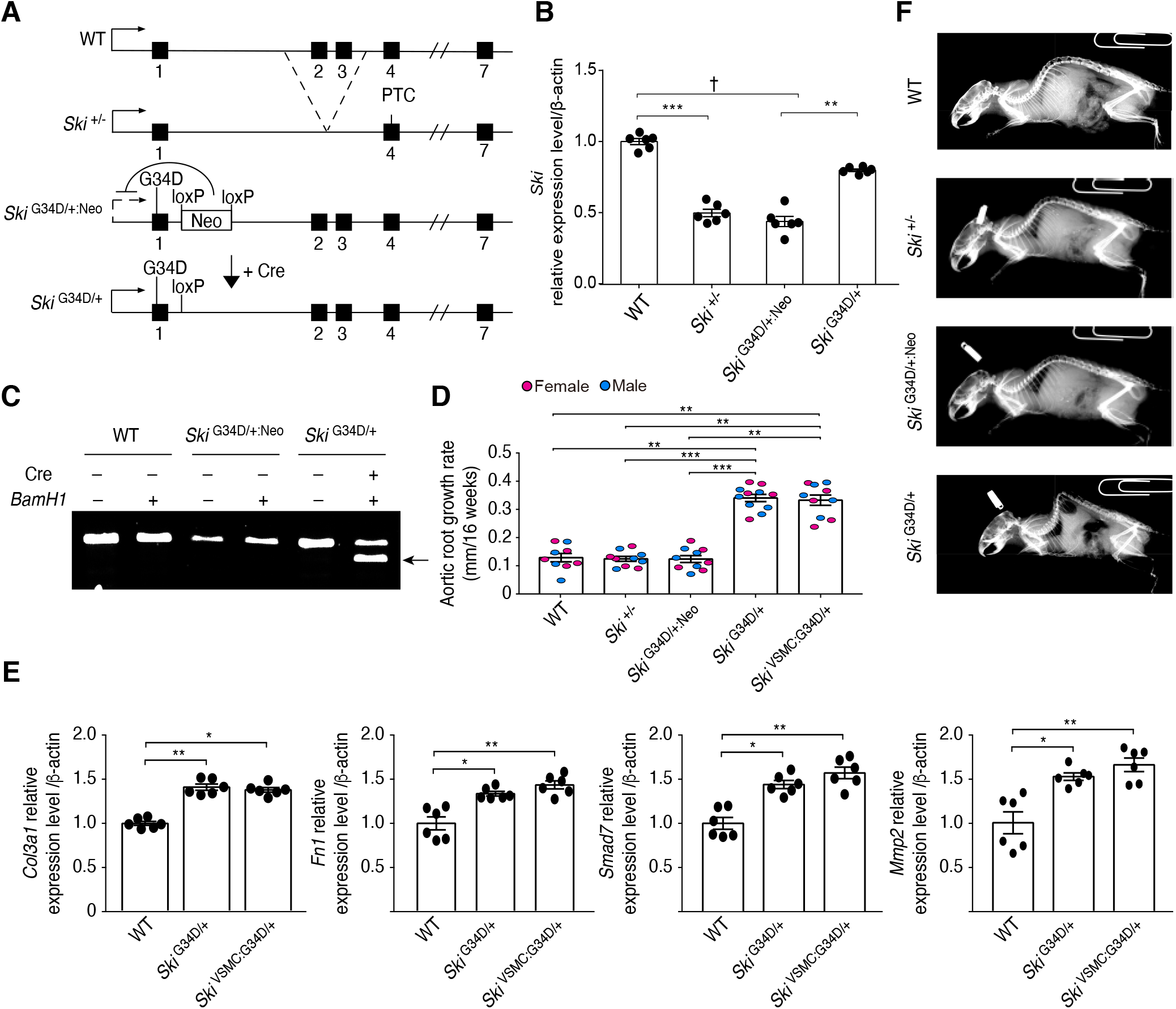
Characterization of SGS mouse models. (A) Structure of *Ski* haploinsufficient and *Ski* mutant alleles in mouse models. Deletion of exons 2 and 3 of *Ski* creates a premature termination codon (PTC) in exon 4 of the *Ski* haploinsufficient allele. (B) Mean expression level (±SEM) of *Ski* mRNA in the tail of each mouse line by qPCR. Both *Ski*^+/−^ *and Ski*^G34D/+:Neo^ haploinsufficient lines express half the normal complement of *Ski* mRNA, while the *Ski*^G34D/+^ mouse line expresses levels comparable to controls; n=6 per group. (C) Expression of WT and mutant *Ski* in each mouse line. The *Ski*^G34D/+^ mouse line uniquely expresses the mutant allele, as shown by the BamH1 restriction fragment of amplified cDNA. (D) Mean aortic root growth rate (±SEM) from 8 to 24 weeks of age. Note that *Ski*^G34D/+^ and *Ski*^VSMC:G34D/+^ mice had significantly greater aortic growth compared to other groups; wild type (n=9), *Ski*^+/−^ (n=10), *Ski*^G34D/+:-Neo^ (n=10), *Ski*^G34D/+^ (n=11), *Ski*^VSMC:G34D/+^ (n=10). (F) Skeletal phenotype: representative spine radio-graphs for each mouse line at 24 weeks of age. Compared to other mouse lines, only *Ski*^G34D/+^ mice demonstrated skeletal deformities. (G) Expression levels of TGFβ target genes (*Col3a1, Fn1, Smad7* and *Mmp2*) relative to β-actin control and normalized to WT expression (±SEM), as determined by qPCR. Compared with WT littermates, *Ski*^G34D/+^ mice and *Ski*^VSMC:G34D/+^ mice demonstrated increased target gene expression; n=6 for all groups. Non-parametric Kruskal-Wallis test with Dunn’s multiple comparison test was used to assess for statistical significance between comparing groups. For all graphs, each bar defines the median with standard error indicated by whiskers and numerical data are presented as scatter dot-plots. *P < 0.05; **P < 0.01; ***P < 0.001; †P < 10^−4^.

Mice heterozygous for the mutant allele (*Ski*^G34D/+^) recapitulated multiple phenotypic characteristics of patients with SGS. In vivo echocardiography of these mice showed evidence of increased aortic root size at 6 months of age and enhanced post-natal aortic root growth from 2 to 6 months of age, compared to wild-type (WT) littermates (Fig. 1D, S1A). They also showed evidence of skeletal deformity in the form of spine kyphosis (Fig. 1F), as well as reduced subcutaneous fat and increased collagen deposition in the skin (Fig. S1B, C), at 6 months of age, in comparison to WT littermates. Finally, they displayed abnormal behavior including hypoactivity and impaired motor performance at 10 weeks of age, when compared to WT littermates (Fig. S1D, E). In contrast, mice haploinsufficient for *Ski (Ski*^+/−^ or *Ski*^G34D/+:Neo^) showed no evidence of these phenotypic defects when compared to WT littermates.

We bred our conditional G34D allele (*Ski*^G34D/+:Neo^) mouse line to transgenic mice carrying the vascular smooth muscle cell (VSMC) specific Sm22α-Cre driver (32), to generate mice that selectively expressed mutant SKI only in VSMC populations. These mice (*Ski*^VSMC:G34D/+^) also showed evidence of aortic root aneurysm at 6 months of age compared to WT littermates, and increased post-natal aortic root growth between 2 and 6 months of age, that was indistinguishable from the rate of growth seen in *Ski*^G34D/+^ mice, which ubiquitously express mutant Ski (Fig. 1D, S1A).

As previously observed in dermal fibroblasts from patients with SGS (21), qPCR of aortic tissue taken from *Ski*^G34D/+^ and *Ski*^VSMC:G34D/+^ mice showed increased expression of all assayed TGFβ target genes, including *Col3A1, Fn1, Smad7, Mmp2, Col1A1, Cdkn1a, Ctgf, Serpine1, Skil, and Mmp9* (encoding type 3 collagen, fibronectin, SMAD family member 7, matrix metallopeptidase 2, type 1 collagen, p21, connective tissue growth factor, PAI-1, SKI-like protein, and matrix metallopeptidase 9, respectively; Fig. 1E, S2).

These data confirm that this knock-in mouse model of SGS recapitulates multiple phenotypic manifestations of the disorder seen in humans, and that the aortic root aneurysm seen in these mice associates with increased TGFβ-dependent target gene expression in the aortic wall. The absence of a phenotype in *Ski*^G34D/+:Neo^ mice supports the conclusion that the SGS phenotype does not manifest as a result of *Ski* haploinsufficiency, leaving open the possibility of either a dominant negative mechanism of action or a novel gain of function.

### Angiotensin-2 type 1 receptor blocker (ARB) losartan ameliorates aortic aneurysm growth in SGS mice

The ARB losartan has previously been shown to ameliorate aortic root aneurysm progression in mouse models of both MFS and LDS, in association with blunted Smad2/3 and Erk1/2 activation (8, 9). To investigate the potential therapeutic effect of losartan in SGS, 2-month old *Ski*^G34D/+^ and *Ski*^VSMC:G34D/+^ mice were treated for 4 months with a dose of losartan previously shown to be efficacious in MFS and LDS mice. Aortic root size was measured at 2 months of age (pre-treatment baseline) and every month thereafter until 6 months of age. Aortic root growth was significantly greater in placebo-treated *Ski*^G34D/+^ and *Ski*^VSMC:G34D/+^ mice compared with WT littermates (Fig. 2A, S3A).

**Figure 2.**
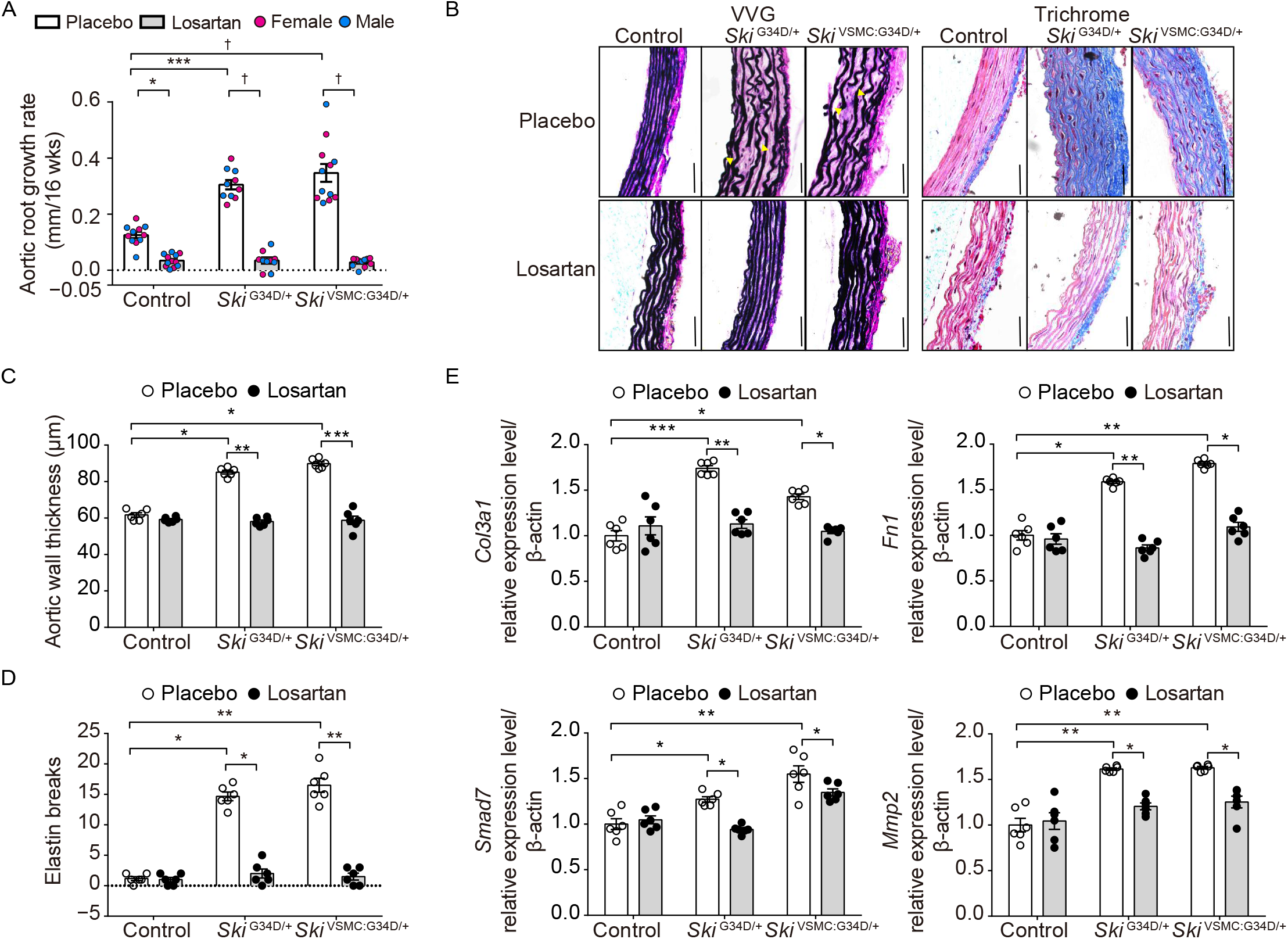
Therapeutic effect of losartan on aortic aneurysm in SGS mice. (A) Mean aortic root growth (±SEM) over 16 weeks of treatment with losartan. Compared with placebo-treated *Ski*^G34D/+^ and *Ski*^VSMC:G34D/+^ mice, losartan-treated *Ski*^G34D/+^ and *Ski*^VSMC:G34D/+^ mice demonstrated reduced aortic root growth. Control–placebo (n=11), *Ski*^G34D/+^–placebo (n=10), *Ski*^VSMC:G34D/+^–placebo (n=12); Control–losartan (n=12), *Ski*^G34D/+^–losartan (n=10), *Ski*^VSMC:G34D/+^ mice–losartan (n=11). (B) Representative cross-sections of the aortic root, stained with VVG (left) and Masson’s trichrome (right), in placebo- and losartan-treated SGS mice; yellow arrow indicates elastic fiber fragmentation. Black line indicates scale bar (50µm). (C) Aortic wall thickness (±SEM). Compared with controls, placebo-treated *Ski*^G34D/+^ and *Ski*^VSMC:G34D/+^ mice demonstrated greater aortic wall thickening, which was significantly reduced in losartan-treated animals; n=6 in all groups. (D) Extent of aortic wall elastic fiber damage, measured by number of elastin breaks (±SEM). Compared with controls, placebo-treated *Ski*^G34D/+^ and *Ski*^VSMC:G34D/+^ mice demonstrated increased elastin breaks, which was significantly reduced in losartan-treated animals; n=6 per group. (E) TGFβ target gene expression relative to β-actin and normalized to control expression (±SEM). Compared with controls, *Ski*^G34D/+^ and *Ski*^VSMC:G34D/+^ mice demonstrated increased TGFβ target gene expression, which was significantly reduced in losartan-treated animals; n=6 per group. Non-parametric Kruskal-Wallis test with Dunn’s multiple comparison test was used to assess for statistical significance between comparing groups. For all graphs, each bar defines the median with standard error indicated by whiskers and numerical data are presented as scatter dot-plots. *P < 0.05; **P < 0.01; ***P < 0.001; †P < 10^−4^.

Compared to placebo-treated *Ski*^G34D/+^ and *Ski*^VSMC:G34D/+^ littermates, aortic root growth was significant reduced in losartan-treated *Ski*^G34D/+^ and *Ski*^VSMC:G34D/+^ mice to a rate indistinguishable from that observed in control mice. No significant change in total body weight was observed with losartan treatment (Fig. S3B). ARBs such as losartan lower blood pressure, which is known to be beneficial in slowing aortic aneurysm growth. To investigate whether losartan was achieving its protective effect solely through blood pressure reduction, we performed a head-to-head trial with another antihypertensive agent, the beta-blocker atenolol. *Ski*^G34D/+^ and *Ski*^VSMC:G34D/+^ mice and control littermates were treated with hemodynamically-equivalent doses of either atenolol (60mg/kg/day) or losartan (50mg/kg/day), from 2 to 6 months of age. Aortic root growth during this 4 month period in atenolol-treated *Ski*^VSMC:G34D/+^ mice was significantly less than that of placebo-treated *Ski*^VSMC:G34D/+^ mice (Fig. S3C), as expected following blood pressure reduction. By contrast, losartan-treated *Ski*^VSMC:G34D/+^ mice showed a significantly greater reduction in aortic root growth when compared to atenolol-treated *Ski*^VSMC:G34D/+^ mice, despite an equivalent reduction in blood pressure and no significant change in total body weight with the 2 drugs (Fig. S3C, D, E)

Histological and morphometric analyses of aortic wall cross sections were performed following death or sacrifice of the mice at 6 months of age. Verhoeff-Van Gieson (VVG) and trichrome staining displayed evidence of aortic wall thickening due to massive accumulation of collagen in the aortic media, reduced elastin content, and increased elastic fiber fragmentation in placebo-treated *Ski*^G34D/+^ and *Ski*^VSMC:G34D/+^ mice, compared with WT littermates, all of which were normalized by losartan treatment (Fig. 2B, C, D).

To further confirm that losartan was achieving its effect through a mechanism other than simple blood pressure reduction, we assessed TGFβ-dependent target gene expression in the aortas of 6-month old mice. Placebo-treated *Ski*^G34D/+^ mice and *Ski*^VSMC:G34D/+^ mice showed a significantly greater expression of TGFβ-dependent target genes, compared with WT littermates (Fig. 2E, S2). By contrast, losartan treatment led to a significant reduction in the expression of these genes in *Ski*^G34D/+^ mice and *Ski*^VSMC:G34D/+^ mice, to levels indistinguishable from those seen in WT littermates in most instances. These data support the conclusion that ARB blockade with the use of losartan appears highly efficacious in treating aortic aneurysm growth in knock-in mouse models of SGS, in association with reduced TGFβ-dependent gene expression, analogous to what has been observed in mouse models of both MFS and LDS (8, 9). It may hence represent a novel therapeutic strategy for the treatment of patients with SGS.

### Selective CBP/P300 inhibition rescues aortic aneurysm growth in SGS mice

The G34D mutation is located in the R-SMAD binding domain of SKI. Interestingly, a mutated form of SKI lacking this R-SMAD binding domain was found to retain its ability to regulate SMAD-mediated transcriptional activation in a transient transfection reporter assay, but failed to dissociate CBP/P300 from the SMAD complex (28, 30). Indeed, SKI and CBP/P300 are known to compete for binding to R-SMADs, and binding of SKI to R-SMADs is sufficient to displace CBP/P300, an effect that is not mimicked by the interaction of SKI with SMAD4. Maintenance of CBP/P300 within the complex promotes gene expression via increased H3K27 acetylation and hence preservation of an open chromatin state (33-39). An increase is resident CBP/P300 can also positively regulate transactivation activity through acetylation of the MH2 domain of SMAD3 at lysine 378 (40). Our prior phenotyping data confirmed that haploinsufficiency for SKI does not appear to be the mechanism of action in SGS. We therefore hypothesized that SGS-causing missense mutations in the R-SMAD binding domain of SKI might allow maintenance of CBP/P300 binding despite residual recruitment of SKI to regulatory elements in target genes via interaction with SMAD4. The resulting increase in H3K27 acetylation and heightened and/or prolonged TGFβ-dependent gene transcription would not manifest in experimental systems that are not susceptible to this type of epigenetic regulation.

To investigate this, we first performed immunofluorescence staining to look for evidence of enhanced H3K27 acetylation in the aortic root of *Ski*^VSMC:G34D/+^ mice at 6 months of age. Compared with control littermates, *Ski*^VSMC:G34D/+^ mice did indeed show much greater H3K27 acetylation in the medial layer of the aortic root (Fig. 3A). To confirm that increased H3K27 acetylation is a driver of aortic aneurysm progression in SGS, rather than simply a marker of it, we treated *Ski*^VSMC:G34D/+^ mice with the selective CBP/P300 inhibitor C646 (41). Treatment with C646 (1 mg/kg/day) was started at 2 months of age and continued for 3 months until sacrifice of the mice at 5 months of age. Control mice were treated with the vehicle dimethyl sulfoxide (DMSO) alone. Aortic root size was measured at 2 months of age (pre-treatment baseline) and every month thereafter. Aortic root growth during the treatment period was significantly greater in vehicle-treated *Ski*^VSMC:G34D/+^ mice, compared with control littermates (Fig. 4A, S4A), and was comparable to that previously seen in placebo-treated *Ski*^VSMC:G34D/+^ mice. By contrast, C646 treatment led to a significant reduction in aortic root growth in *Ski*^VSMC:G34D/+^ mice, to a rate indistinguishable from that observed in control mice, without a change in total body weight (Fig. S4B). Importantly, C646 therapy had no effect in control mice, showing that inhibition of H3K27 acetylation was selectively targeting *Ski* mutation-associated pathological aortic root growth rather than physiological aortic growth.

**Figure 3.**
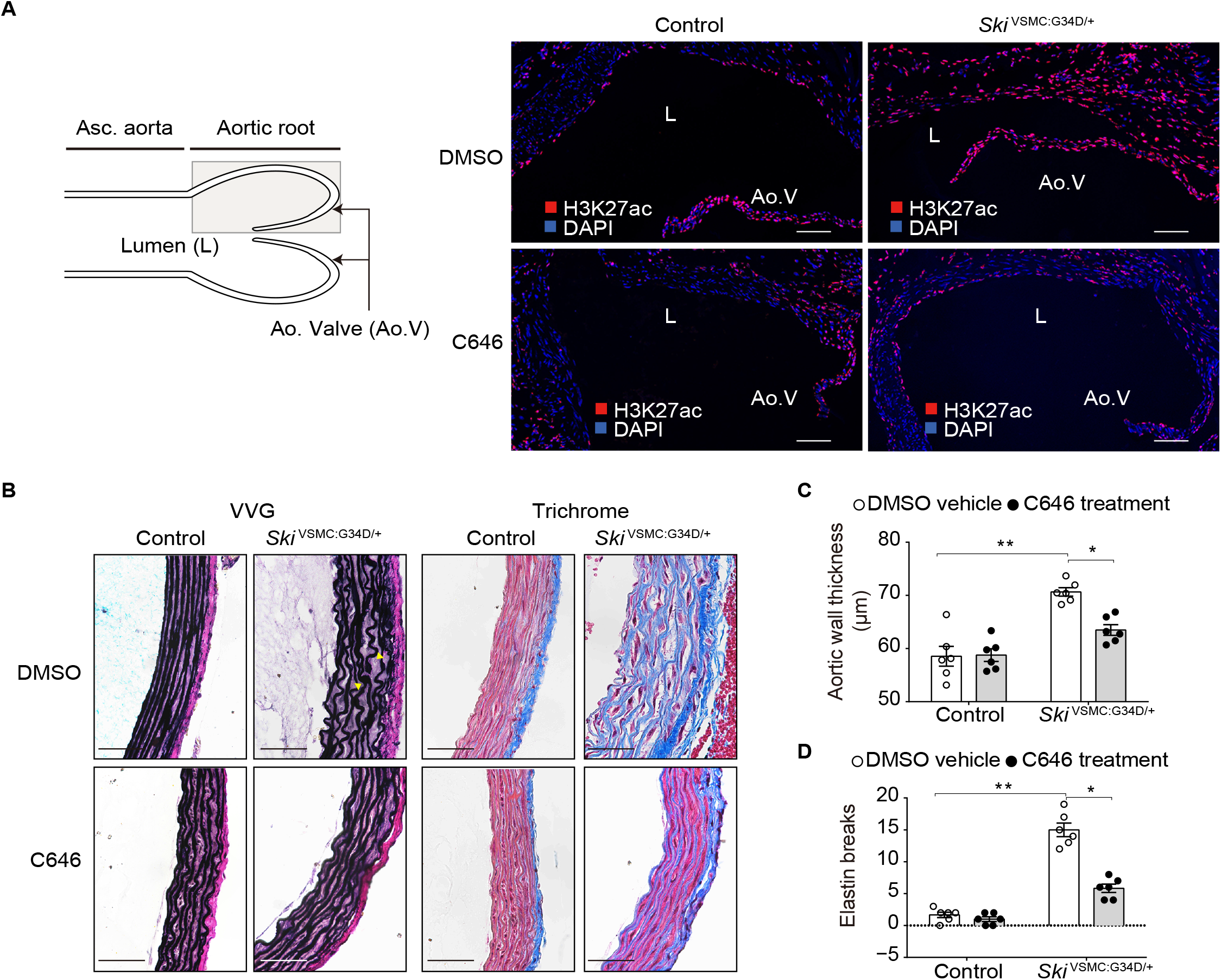
Phenotypic effect of CBP/P300 inhibition using C646 on aortic root aneurysm in mouse models of SGS. (A) Representative immunofluorescence image of aortic root samples of control and *Ski*^VSMC:G34D/+^ mice treated with either C646 or vehicle (DMSO) probed for histone 3 lysine 27 acetylation (H3K27ac). Compared with controls, *Ski*^VSMC:G34D/+^ mice demonstrated increased acetylation, which was reduced by C646 treatment. White line indicates scale bar (50µm). (B) Aortic root histology in DMSO- and C646-treated control and *Ski*^VSMC:G34D/+^ mice. Images show representative cross-sections stained with VVG (left) and Masson’s trichrome stain (right). Yellow arrows indicate elastic fiber breaks. Black line indicates scale bar (50µm). (C) Aortic wall thickness (±SEM) in DMSO- and C646-treated control and SGS mice. Compared with controls, DMSO-treated *Ski*^VSMC:G34D/+^ animals showed greater medial wall thickening, which was reduced in C646-treated *Ski*^VSMC:G34D/+^ littermates; n=6 for all groups. (D) Mean number of elastin breaks (±SEM) in DMSO- and C646-treated control and SGS mice. Compared with controls, DMSO-treated *Ski*^VSMC:G34D/+^ mice showed more elastin breaks, which was prevented by C646 treatment; n=6 for all groups. Non-parametric Kruskal-Wallis test with Dunn’s multiple comparison test was used to assess for statistical significance between comparing groups. For all graphs, each bar defines the median with standard error indicated by whiskers and numerical data are presented as scatter dot-plots. *P < 0.05; **P < 0.01.

**Figure 4.**
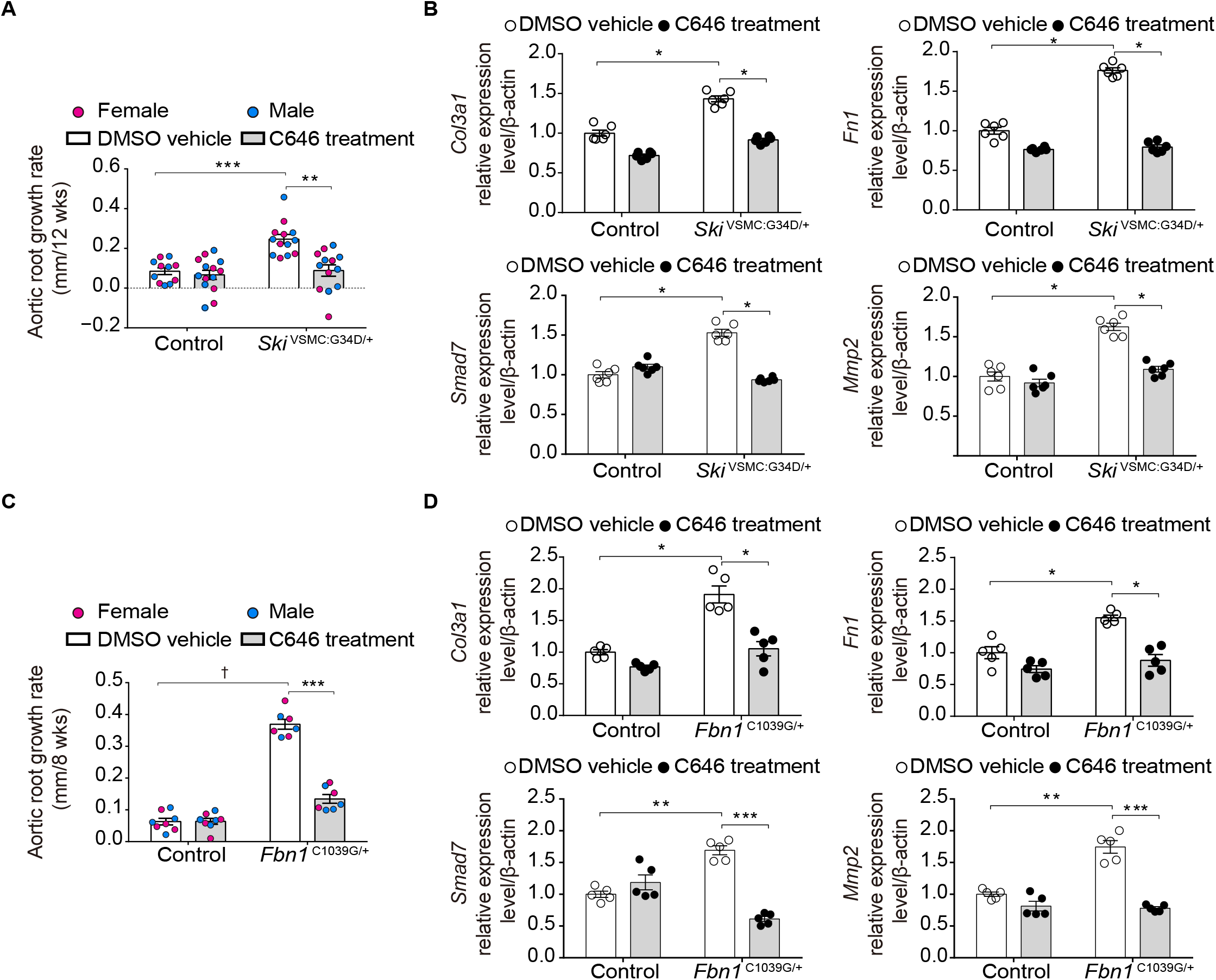
Therapeutic effect of CBP/P300 inhibition using C646 on aortic root aneurysm in mouse models of SGS and MFS. (A) Mean aortic root growth over 12 weeks of treatment with C646 in control and *Ski*^VSMC:G34D/+^ mice. Compared with controls, *Ski*^VSMC:G34D/+^ mice demonstrated significantly greater aortic root growth, which was reduced by treatment with C646; Control–DMSO (n=11), *Ski*^VSMC:G34D/+^–DMSO (n=13); Control–C646 (n=14), *Ski*^VSMC:G34D/+^–C646 (n=13). (B) TGFβ target gene expression relative to β-actin and normalized to control expression (±SEM), as determined by qPCR. Compared with controls, *Ski*^VSMC:G34D/+^ mice showed increased target gene expression, which was reduced by C646 treatment; n=6 per group. (C) Mean aortic root growth over 8 weeks of treatment with C646 in control and MFS (*Fbn1*^C103934G/+^) mice. Compared with DMSO-treated MFS mice, C646-treated MFS animals had a significant reduction in aortic root growth rate; Control–DMSO (n=8), MFS–DMSO (n=8); Control–C646 (n=7), MFS–C646 (n=7). (D) TGFβ target gene expression relative to β-actin and normalized to control expression (±SEM), as determined by qPCR. Compared with controls, MFS (*Fbn1*^C103934G/+^) mice had significantly increased target gene expression, which was reduced in C646-treated MFS animals. Non-parametric Kruskal-Wallis test with Dunn’s multiple comparison test was used to assess for statistical significance between comparing groups. For all graphs, each bar defines the median with standard error indicated by whiskers and numerical data are presented as scatter dot-plots. *P < 0.05; **P < 0.01; ***P < 0.001; †P < 10^−4^.

The specificity of C646 was confirmed by immunofluorescence staining for H3K27 acetylation. Compared with DMSO-treated *Ski*^VSMC:G34D/+^ littermates, C646-treated *Ski*^VSMC:G34D/+^ mice showed a clear reduction in H3K27 acetylation in the medial layer of the aortic root (Fig. 3A). Histological and morphometric analyses of aortic wall cross sections were performed following death or sacrifice of the mice at 5 months of age. VVG and trichrome staining reconfirmed prominent aortic wall thickening, reduced elastin content, and increased elastic fiber fragmentation in vehicle-treated *Ski*^VSMC:G34D/+^ mice compared to control littermates, all of which were significantly improved in C646-treated *Ski*^VSMC:G34D/+^ mice (Fig. 3B, C, D).

To determine whether this reduction in H3K27 acetylation by C646 directly impacted TGFβ-dependent gene expression in the aortas of these mice, we performed qPCR on the aortas of 5-month old mice. The increased expression of TGFβ target genes seen in vehicle-treated *Ski*^VSMC:G34D/+^ mice was indeed significantly reduced in C646-treated *Ski*^VSMC:G34D/+^ mice, to levels close to or indistinguishable from those seen in control mice (Fig. 4B, S5). To confirm that C646 does not have any antihypertensive effect, we measured blood pressure in these animals and found no significant difference between vehicle-treated *Ski*^VSMC:G34D/+^ mice and C646-treated *Ski*^VSMC:G34D/+^ littermates (Fig. S4C).

These data suggest that missense mutations in the R-SMAD domain of SKI lead to enhanced CBP/P300 activity, increased H3K27 acetylation, and ultimately prolonged TGFβ-dependent gene expression, all of which can be ameliorated in a mouse model of SGS by selective CBP/P300 inhibition. Since animal models of MFS, in particular the *Fbn1*^C1039G/+^ mouse model, also show increased TGFβ-dependent gene expression in their aortas, we hypothesized that selective CBP/P300 inhibition could represent a novel therapeutic strategy for other forms of inherited aortic aneurysm. To assess this, we treated *Fbn1*^C1039G/+^ mice with C646 from 2 months of age for 2 months. Aortic root growth over the treatment period was significantly greater in vehicle-treated *Fbn1*^C1039G/+^ mice than WT littermates (Fig. 4C). By contrast, C646 treatment led to a significant reduction in aortic root growth in *Fbn1*^C1039G/+^ mice, to a rate comparable to that observed in WT controls. This reduction in aortic root growth was associated with a significant reduction in TGFβ-dependent gene expression in C646-treated *Fbn1*^C1039G/+^ mice, compared to their vehicle-treated littermates (Fig. 4D).

### CBP/P300 inhibition abrogates TGFβ-dependent gene expression in SGS patient cells

We tested the effect of C646 on primary dermal fibroblasts derived from 2 SGS patients and 2 healthy controls. Vehicle-treated fibroblasts from SGS patients showed significantly greater mRNA expression of TGFβ-dependent genes, including *COL3A1, FN1, SKI*, and *SMAD7*, compared with control fibroblasts, both at baseline and after stimulation with exogenous TGFβ1 (10 ng/ml) for 6 hours (Fig. S6). By contrast, C646 treatment of SGS patient fibroblasts led to a significant reduction in expression of these genes, to levels similar to those seen in control fibroblasts, or SGS patient fibroblasts treated with the TGFβ type I receptor (Alk5) kinase inhibitor SD208. Selective CBP/P300 inhibition thus appears to suppress TGFβ-dependent gene expression in both mouse models of SGS and cells derived from patients with SGS.

## Discussion

Shprintzen-Goldberg syndrome (SGS) in a rare systemic connective tissue disorder caused by heterozygous mutations in the *SKI* gene, and shows significant phenotypic overlap with both MFS and LDS. Despite the initial discovery of a causal gene, a number of questions remained unanswered about the disorder, including the effect of SGS-causing mutations on TGFβ signaling in vivo, the exact role of TGFβ signaling in the disease, and which SKI binding partners and downstream signaling sequelae contribute to pathogenesis.

The body of evidence implicating dysregulation of TGFβ signaling in vascular connective tissue disorders is extensive and compelling. Virtually every study of these conditions has provided evidence for high TGFβ signaling in the aortic wall of mouse models or people with MFS or LDS, including enhanced activation of signaling intermediates (i.e. phosphorylation and/or nuclear translocation of SMAD2/3) and high output of TGFβ target genes in relevant tissues. Yet ambiguity is demonstrable and controversy substantial. Evidence suggested that fibrillin-1, the deficient gene product in Marfan syndrome, could positively regulate TGFβ signaling by concentrating cytokine at sites of intended function, but negatively regulate signaling by sequestering the TGFβ latent complex from activators (5, 42-44). All forms of LDS are caused by heterozygous loss-of-function mutations in genes encoding positive effectors of TGFβ signaling (4, 17, 45-47). Therapeutic trials were equally ambiguous. While the consistent therapeutic benefit of ARBs in mouse models of MFS or LDS strictly correlated with attenuation of the tissue signature for high TGFβ signaling in the aortic wall, administration of TGFβ neutralizing antibodies accentuated disease in the perinatal period in MFS mice, while affording significant protection later in postnatal life, with heightened efficacy when used in combination with ARBs (9). The fact SGS includes essentially every systemic manifestation of MFS (except lens dislocation) and LDS seemed particularly relevant given the prominent role of SKI in the negative regulation of the TGFβ transcriptional response (1, 2, 21, 48). The observations that SGS mutations clustered in the R-SMAD binding domain of SKI, and that SGS patient fibroblasts showed high expression of TGFβ-responsive genes known to contribute to aneurysm progression suggested that increased TGFβ-dependent events may cause the multisystem manifestations of SGS. This might also inform the mechanism for similar manifestations in MFS and LDS.

More recently, it was reported by Hill and colleagues that SGS mutations can promote enhanced stability of SKI, and that this was associated with decreased expression of selected TGFβ target genes in either cells transfected with mutant SKI or in SGS fibroblasts (31). Notably, this study only assessed the acute phase-response to administered TGFβ ligand (1 and 8 hours after delivery), and focused on genes that are predominantly expressed in neurons or polarized epithelial cells, but not the aortic wall or dermal fibroblasts. Prior work had demonstrated that loss of binding of SKI to R-SMADs, as imposed by SGS mutations, did not abrogate the regulation of the TGFβ transcriptional response in reporter allele assays, but rather impaired the ability of SKI to displace CBP/P300 at critical regulatory elements, potentially altering the efficiency of chronic SKI-mediated termination of a signal initiated by TGFβ (28, 30). This effect would be best interrogated upon chronic exposure to TGFβ, with potentially unique insights afforded by the study of affected tissues in vivo. Moreover, the impact of an SGS mutation could vary based upon cell type, with particular relevance for the expression of redundant negative regulators (e.g. SKIL or SMAD7) or factors involved in chronic compensation. Importantly, our prior studies in dermal fibroblasts derived from SGS patients assessed TGFβ target gene expression at steady-state, with no overlap between the repertoire of genes previously assessed and those specifically examined by Hill and colleagues (21, 31). We now unequivocally show that constitutive or VSMC-specific expression of a heterozygous SGS-associated SKI mutation leads to a substantial and sustained increase in H3K27 acetylation in vivo, as predicted by increased CBP/P300 occupancy, in association with high expression of TGFβ target genes relevant to aortic disease and reliably assayed in dermal fibroblasts. These findings correlated with the excessive accumulation of fibrillar collagens in the vessel wall and skin of SGS mouse models, as predicted by amplification of the TGFβ transcriptional response, with notable downregulation appearing coincident with therapeutic interventions that achieved attenuation of aneurysm progression.

These data are consistent with prior work showing that CBP/P300-mediated H3K27 acetylation is enriched in the promoter regions of TGFβ target genes (49). Furthermore, CBP/P300-mediated histone acetylation at the PAI-1 and p21 promoters can enhance TGFβ1-induced expression of these genes in cell culture systems (50). By contrast, CBP/P300 inhibition using C646 has been shown to significantly reduce H3K27 acetylation (51), and also abrogate expression of a number of TGFβ target genes, including *Cdkn1a, Mmp, Serpine1* and *Ctgf* (52-55). Prior work has shown that C646 can have a substantial effect on behavioral characteristics in rodents such fear memory (56, 57), and on cardiac fibrosis and hypertrophy in *Sirt3* deficient mice (58). Although we have shown efficacy of C646 against aortic aneurysm progression in SGS and MFS mice, its therapeutic potential for other systemic manifestations of these vascular connective tissue disorders remains untested.

ARBs such as losartan have previously been shown to attenuate TGFβ signaling through downregulation of TGFβ ligands, receptors and activators such as PAI-1 (9, 11). The overt protection from aneurysm progression seen in mouse models of MFS and LDS associates with a reduction in TGFβ signaling in the aortic root media, as evidenced by normalization of SMAD2/3 phosphorylation and the expression of TGFβ target genes in the aortic wall, prominently including fibrillar collagens, MMPs 2 and 9 and CTGF (7, 8, 11). A role for losartan in epigenetic modulation of the TGFβ transcriptional response is less clear. Work by Reddy et al. has shown that losartan can ameliorate diabetic nephropathy in mice through a reduction in H3K9/14 acetylation at the promoters of pathogenic genes (59). Furthermore, losartan was found to attenuate proteinuric kidney disease in mice via inhibition of DNA methylation at the nephrin promoter (60). The relative importance of each of these potential mechanisms of action for ARBs in SGS remains to be determined. Our current hypothesis is that ARBs predominantly suppress early events in TGFβ signaling, precluding abnormal maintenance and/or amplification of the transcriptional response in the context of a SKI functional deficiency.

Given the known phenotypic overlap, and the newly-confirmed biochemical similarity, between SGS and both MFS and LDS, a logical extension of this work was to elucidate whether epigenetic changes and their pharmacological manipulation may hold relevance to other CTDs. Indeed, treatment with C646 did lead to a significant reduction in aortic root growth in MFS mice, in association with reduced TGFβ target gene expression in the aortic wall. This suggests that a reduction in H3K27 acetylation can achieve a therapeutic effect, likely by facilitating a more closed chromatin state that impairs TGFβ target gene transcription, despite the presence of increased upstream TGFβ signaling. The findings of this study suggest that epigenetic modulation holds potential for the treatment in diverse presentations of syndromic forms of aortic aneurysm.

These data add to the extensive and compelling in vivo evidence for enhanced TGFβ signaling in the pathogenesis of vascular connective tissue disorders including MFS, LDS and SGS, with particular emphasis on the vascular pathology. In comparison, there is no documented example of decreased TGFβ signaling in tissues derived from people or mouse models of these conditions. Yet, we view the evidence for primary functional impairment of the TGFβ signaling response by underlying mutations to be equally compelling, including the evidence by Hill and colleagues that SGS mutations can stabilize SKI and associate with relative impairment of transcriptional responses in selected culture systems (31). The challenge – and, we would argue, the opportunity - lies in embracing all inconvenient truths to arrive at reconciling and testable models that have the true potential to inform disease pathogenesis and therapeutic opportunities. In the case of LDS, we have shown that heterozygous loss-of-function mutations in positive effectors of TGFβ signaling have a disproportionate and at times even unique negative impact on responses in specific cell types (12). This can lead to paracrine effects (e.g enhanced TGFβ ligand production) that drive excessive signaling in neighboring cell types that are less vulnerable to the consequences of underlying mutations (12). Given the potency and redundancy of mechanisms for autoregulation of TGFβ signaling, it seems possible that the apparent low signaling – high signaling paradox in the TGFβ vasculopathies is actually a requirement for the initiation and maintenance of a high TGFβ signaling state. There is ample precedent for this in the TGFβ cancer paradox, where TGFβ serves both as a tumor suppressor and a positive regulator of tumor progression. We anticipate that consideration and integration of this physiologic complexity will be required to achieve consensus – and truth – in our field.

## Methods

### Mouse lines

All mice were cared for under strict compliance with the Animal Care and Use Committee of the Johns Hopkins University School of Medicine. *Ski* ^G34D/+^ mice were generated by homologous recombination, as described in the next section. *Ski* ^tm1a(EUCOMM)Hmgu^ (tm1a represents targeted mutation 1a, and hmgu represents Helmholtz Zentrum Muenchen GmbH) embryonic stem cells were obtained from the European Conditional Mouse Mutagenesis Program and injected into the cavity of day 3.5 blastocysts from C57BL/6J mice at the Johns Hopkins University School of Medicine transgenic core. Male chimeras were mated with C57BL/6J WT female mice to establish germline transmission. The *LacZ*-Neo cassette was removed by crossing with a FlpO transgenic strain (B6.Cg-Tg(Pgk1-flpo)10Sykr/J, #011065) purchased from the Jackson Laboratory and to generate *Ski* ^+/−^ mice, exon 2 and 3 of *Ski* gene flanking by *loxP* sequences were removed by crossing with a transgenic Cre strain (B6.C-Tg(CMV-cre)1Cgn/J, #006054) purchased from the Jackson Laboratory, followed by mating to the C57BL/6J strain at least 5 generations. Sm22α-Cre mice (B6.Cg-Tg(Tagln-cre)1Her/J, #017491) were purchased from the Jackson Laboratory, followed by mating to the C57BL/6J strain for at least 5 generations. To minimize potentially confounding background effects, all comparisons between genotypes and between treatment arms within a genotype were made between gender-matched littermates.

Mice were checked daily for evidence of premature lethality. At the end of a drug trial, all mice were euthanized through inhalational halothane (Sigma) or anesthetized with isofluorane. Following sacrifice, mice underwent immediate laparotomy, descending abdominal aortic transection, and phosphate-buffered saline (PBS; pH 7.4) was infused throughout the vascular tree via the left ventricle. For the aorta frozen section, additional 4% paraformaldehyde in PBS was infused again for fixation tissue. Sacrificed mice used for 3 harvest methods, latex infusion for histological analysis, freezing heart and aorta embedded in optimal cutting temperature compound (O.C.T compound) for immunofluorescent staining, and *in-situ* hybridization and snap-frozen aorta in liquid nitrogen.

For quantitative RT-PCR, the aortic root and ascending aorta (above the aortic root to the origin of right brachiocephalic trunk) of mice were harvested separately, snap-frozen in liquid nitrogen, and stored at −80°C until processed. For RNA extraction, aortas were homogenized in TRizol (ThermoFisher) by FastPrep-24 (MP Biomedicals, LLC), per the manufacturer’s instructions. After homogenization, RNA was extracted using an RNeasy mini kit (QIAGEN), per the manufacturer’s instructions. The RNA samples were then stored once more at −80°C until quantitative RT-PCR was performed.

For the frozen aorta sections, aorta with heart was harvested and fixed in fresh 4% paraformaldehyde in PBS at 4°C overnight, and then placed in cold 30% sucrose in PBS solution and incubated at 4°C overnight again. Tissue was then embedded in Tissue-Tek O.C.T compound and snap-frozen in liquid nitrogen, and stored at −80°C until processed.

According to a previously described protocol with slight modification (7, 11, 61) mice that were analyzed for aortic histology had latex infused into the left ventricle through the descending abdominal aorta. Mice were then fixed for 24 hours in 10% neutral-buffered formalin, before being stored in 70% ethanol until the histological analysis was performed.

### Generation of Ski ^G34D/+^ mice

*Ski* ^G34D/+^ mice were generated by homologous recombination. A 10-kb *Ski* fragment was generated by PCR from mouse embryonic stem cell DNA. The amplicon was subcloned into pCR2.1-TOPO (Invitrogen Corp.). Site-directed mutagenesis was performed with the In-Fusion HD kit (Clontech Inc.), creating G34D mutation. The Neo cassette was amplified from pMC1neo-polyA vector (Stratagene Inc.) and the fragment containing the *Sal*1 restriction site and Neo cassette with flanking *loxP* sequences was subcloned into a unique *Sal*1 site in the *Ski* intron after exon 1. All targeting vector sequences, including the sequences of the *loxP* sites and site-directed mutagenesis-created mutations were confirmed by sanger sequencing. The vector was linearized using a unique (*EcoR*1) site and electroporated into R1 embryonic stem cells. Positive clones were identified by PCR test. Positives clones were injected into 129S6/ScEvTac blastocysts at embryonic day 3.5 and transferred into pseudopregnant females. Chimeric offspring were mated to C57BL/6J mice, and germline transmission was observed for at least 3 independent targeting events for mutant genotype. All exons encompassed by and immediately flanking the targeting vector were analyzed by sequencing of PCR-amplified genomic DNA derived from mutant mice, to demonstrate the fidelity of targeting. Mice were genotyped on the basis of creation of a new *BamH*1 site in correctly targeted mice. Primers used for amplification were 5’–GAGCCCGATCG CACCATGGAA-3’ (sense) and 5’-AAGAGATGGTCTCCCCTTCC-3’ (antisense). For testing the random insertion of linearized targeted vector, quantitative PCR of the Neo cassette sequence was performed with a previously verified DNA sample, which contained only a single Neo cassette sequence. The *loxP*-flanked Neo cassette was removed by crossing with a Cre deleter strain, either CMV-Cre (B6.C-Tg(CMV-cre)1Cgn/J, #006054) purchased from the Jackson Laboratory, followed by mating to C57BL/6J strain for at least 5 generations for a C57BL/6J genetic background, or Prm-Cre (129S/Sv-Tg(Prm-Cre)58Og/J, #003328) purchased from Jackson Laboratory, followed by mating to a 129S6/ScEvTac strain for at least 5 generations for a 129S6/ScEvTac genetic background. The 129S6/ScEvTac genetic background *Ski* ^G34D/+^ mice were used for analyses of dermal, skeletal, and cardiovascular phenotypes. The C57BL/6J genetic background *Ski* ^G34D/+^ mice were used for behavioral phenotype analysis. *Ski* ^G34D/+:Neo^ mice were bred to the 129S6/ScEvTac strain for at least 5 generations, without deletion of the Neo cassette as a separate mouse line. Complete concordance of phenotype for 3 or 2 independent lines and backcrossing to each congenic inbred strain for at least 5 generations excluded any major off-target effects.

### Mouse drug treatment

Losartan was dissolved in drinking water and filtered to reach a concentration of 0.5 g/L, giving an estimated daily dose of 50 mg/kg/day (based on a 30-g mouse drinking 3 mLs per day). Atenolol was dissolved in drinking water and filtered to reach a concentration of 0.6 g/L, giving an estimated daily dose of 60 mg/kg/day. Placebo-treated mice received drinking water. Mice given these medications were started on treatment at 8 weeks of age and continued for 16 weeks. C646 (Selleckchem, #S7152) was reconstituted in 10% DMSO (Sigma) dissolved in PBS, and administered daily by intraperitoneal injection at a dose of 1 mg/kg/day. Treatment was initiated at 8 weeks of age and continued for 12 weeks. Ten percent DMSO in PBS was administered as a control.

### Mouse echocardiography

Nair hair removal cream was used to remove fur from the anterior thorax of the mice the day prior to echocardiography. According to a previous protocol with slight modifications (7, 11, 61), echocardiography was performed on awake, nonsedated mice using a Vevo 2100 imaging system and 40 MHz transducer (Visualsonics). Mice were imaged at 8 weeks of age as a baseline and every 4 weeks thereafter, until 20 weeks of age. The aorta was imaged in the parasternal long-axis view. Three separate measurements of the maximal internal dimension at the sinus of Valsalva during systole were made in separate cardiac cycles and averaged. All imaging and analysis was performed blinded to genotype and treatment arm.

### Mouse blood pressure

According to a previously described protocol with slight modifications (7, 11, 61), blood pressure was measured by tail-cuff plethysmography using a Harvard Apparatus IITC noninvasive tail cuff device. Mice were placed in a standard acrylic restrainer for adult mice, with an internal diameter of 25 mm and an adjustable head gate. The end plate was removable, allowing the mice to walk into the restrainer without using force. Hemodynamic recordings were made without sedation or anesthesia. Blood pressure was measured at the end of the drug trial. Mice were habituated for 4 days. On day 5, ten blood pressures were obtained and averaged.

### Mouse radiography

According to a previously described protocol with slight modifications (7, 11, 61), Mice were anesthetized using a combination of 50 mg/kg of ketamine-HCl and 5 mg/kg xylazine-HCl by intraperitoneal injection before X-ray imaging. Mice were placed in the left lateral decubitus position on a radiolucent platform with a metal paper clip as a scale bar and imaged at 1*x* magnification using a Faxitron MX20 (Faxitron).

### Mouse aorta histological analysis

According to a previously described protocol with slight modifications (7, 11, 61), latex-infused heart and aorta were removed from body and transected just below the level of the aortic annulus, and just above the aortic root, and 2-to 3-mm transverse sections were mounted in 4% agar prior to fixation in paraffin. Five-micron aortic sections underwent Verhoeff-van Giesen (VVG) and Masson’s trichrome staining and were imaged at 40*x* magnification using a Nikon Eclipse E400 microscope. Aortic wall thickness was measured of 4 sites of 4 representative sections for each mouse. The disruptions of elastin fiber architecture were counted in 4 sections every 25 microns from the aortic annulus. All analyses were performed blinded to genotype and treatment arm and the results were averaged.

### Mouse behavioral tests

Behavioral tests were performed per the manufacturer’s instructions. All of behavioral tests were performed at 10 weeks of age. For the open field test, a mouse was placed into a corner of a 45 x 45 cm open-filed chamber (San Diego Instruments) with a 16 x 16 photobeam configuration. The behavior of the mouse was monitored for 5 minutes in each of 6 cycles (total 30 minutes). Mouse activity in center area and/or peripheral areas was recorded by beam interruptions. Total activity was calculated by adding up all beam interruptions during the cycle. Activity in the central and peripheral areas was calculated by adding up the beam interruptions in these two areas, respectively. For the rotarod test, mice were placed on a horizontally oriented, rotating cylinder (rod) suspended above the cage floor (Rotamex-5 rotarod, Columbus Instruments). The acceleration started at 0 rpm and was increased by 1.0 rpm every 5 seconds. Velocity and time were recorded at the time of falling for 3 measurements. Each mouse was given 3 minutes of rest time between trials. All analyses were performed blinded to genotype and the results were averaged.

### Human cell culture

Primary human dermal fibroblasts were derived from forearm skin biopsies of 2 controls and 2 patients with SGS, as previously described in (21). The fibroblasts were cultured in Dulbecco’s modified eagle medium (DMEM) with 10% fetal bovine serum (FBS) in the presence of antibiotics and passaged confluence. According to a previously described protocol with slight modifications (21), all cell culture experiments were conducted in serum-starved media for 24 hours prior to drug treatment. Stimulation was performed using 10 ng/ml recombinant human TGFβ1 (R&D system). C646 dissolved in DMSO was treated at a dose of 20 μM for 24 hours of pretreatment before TGFβ1 stimulation. SD208 dissolved in DMSO was treated at a dose of 10 μM for 24 hours of pretreatment before TGFβ1 stimulation. Cells were collected at baseline, 6 hours after TGFβ1 stimulation. The RNA was extracted from the cells using TRizol (ThermoFisher) via an RNeasy mini kit (QIAGEN), per the manufacturer’s instructions.

### Quantitative RT-PCR expression analysis

Total RNA was isolated from mouse aortas or cultured cells using an RNeasy mini kit (QIAGEN), per the manufacturer’s instructions. Quantitative PCR was performed in triplicate with TaqMan Universal PCR Master Mix using an ABI Prism 7900 HT QPCR machine (all from Applied Biosystems), per the manufacturer’s instructions. The following prevalidated TaqMan probes were used to detect specific transcripts: Mm00801666_g1 (*Col1a1*), Mm01254476_m1 (*Col3A1*), Mm01256744_m1 (*Fn1*), Mm04205640_g1 (*Cdkn1a*), Mm01192932_g1 (*Ctgf*), and Mm00435860_m1 (*Serpine1*), Mm00448744_m1 (*Ski)*, Mm00456917_m1 (*Skil*), Mm00484742_m1 (*Smad7*) and Mm00607939_s1 (*Actb*) (Life Technologies). For human samples, the following probes were used: Hs00943809_m1 (*COL3A1*), Hs00365052_m1 (*FN1*), Hs00161707_m1 (*SKI*), Hs01045418_m1 (*SKIL*) and Hs01060665_g1 (*ACTB*). Reactions were run in triplicate, and relative quantification for each transcript was obtained by normalization against a housekeeping control transcript, such as *β–ACTIN* (*ACTB*), according to the formula 2^-Ct^/2^-Ct(ACTB)^.

### Immunofluorescence

Immunofluorescence was performed as previous described (8, 9). Frozen 10-μM long-axis– view sections were obtained with a cryostat and mounted on glass slides. Sections were dried at room temperature overnight prior to staining. Sections were permeabilized in staining buffer (PBS containing 0.1% Triton-× 100) for 30 minutes and then incubated with Fc Receptor Block from Innovex for 30 minutes at room temperature, washed briefly in staining buffer, and then incubated again in blocking solution (0.1% Triton-× 100, 1:50 goat serum, 0.3M glycine) for 30 minutes. Primary antibodies were diluted at 1:100 in staining buffer and incubated with goat anti-rabbit secondary antibody conjugated to Alexa Fluor 555 (Life Technologies) at 1:200 for 1 hour before being mounted with VECTASHIELD Hard Set Mounting Medium with DAPI. Images were acquired on a Zeiss Axio Examiner with a 710NLO-Meta multiphoton confocal microscope at 25*x* magnification. The following primary antibody was used: anti-H3K27ac (Abcam, ab4729).

### Statistics

All quantitative data are shown as bar graphs, produced using Graphpad Prism. Mean ± standard errors of the mean (SEM) are displayed. Statistical analyses were performed using non-parametric test (Kruskal-Wallis test with Dunn’s multiple comparison test). A *P* value < 0.05 was considered to be statistically significant for all tests.

### Study approval

This study was performed in accordance with the recommendations in the Guide for the Care and Use of Laboratory Animals of the National Institutes of Health. All of the animals were handled according to approved institutional animal care and use committee (IACUC) protocols of the Johns Hopkins University School of Medicine. The protocol was approved by the Committee on the Ethics of Animal Experiments of the Johns Hopkins University School of Medicine.

## Author contributions

H.C.D and B.E.K. developed the concept. B.E.K. generated mouse models, designed, performed and directed experiments, analyzed data and wrote the manuscript. J.J.D., E.G.M., and H.C.D. aided in experimental design and interpretation of the data. R.B. aided in mouse colony maintenance and drug treatment. D.B. aided in mouse echocardiography and analyses. R.B., J.J.D., E.G.M., and H.C.D provided essential expertise in the editing of the manuscript. All authors discussed the results and commented on the manuscript prior to submission.

## Acknowledgements

This work was supported by the Howard Hughes Medical Institute, the Johns Hopkins University School of Medicine Cellular and Molecular Medicine graduate training program (T32GM008752). We thank the Johns Hopkins University School of Medicine Transgenic Mouse Core laboratory for expert technical assistance with generating mouse models; J. Habashi for echocardiography analysis; the Johns Hopkins Hospital Department of Pathology Reference laboratory for histology; the Johns Hopkins University School of Medicine Behavioral Core laboratory for mouse behavioral testing.

## References

1. Schepers D et al. The SMAD-binding domain of SKI: a hotspot for de novo mutations causing Shprintzen&ndash;Goldberg syndrome. Eur J Hum Genet 2014;23(2):1–5.

2. Pagon RA et al. Shprintzen-Goldberg Syndrome 1993;

3. Dietz HC et al. Marfan syndrome caused by a recurrent de novo missense mutation in the fibrillin gene. Nature 1991;352(6333):337–339.

4. Loeys BL et al. A syndrome of altered cardiovascular, craniofacial, neurocognitive and skeletal development caused by mutations in TGFBR1 or TGFBR2. Nat Genet 2005;37(3):275– 281.

5. Neptune ER et al. Dysregulation of TGF-beta activation contributes to pathogenesis in Marfan syndrome. Nat Genet 2003;33(3):407–411.

6. Judge DP et al. Evidence for a critical contribution of haploinsufficiency in the complex pathogenesis of Marfan syndrome. J. Clin. Invest. 2004;114(2):172–181.

7. Holm TM et al. Noncanonical TGF Signaling Contributes to Aortic Aneurysm Progression in Marfan Syndrome Mice. Science 2011;332(6027):358–361.

8. Gallo EM et al. Angiotensin II-dependent TGF-β signaling contributes to Loeys-Dietz syndrome vascular pathogenesis. J. Clin. Invest. 2014;124(1):448–460.

9. Habashi JP et al. Losartan, an AT1 Antagonist, Prevents Aortic Aneurysm in a Mouse Model of Marfan Syndrome. Science 2006;312(5770):117–121.

10. Cohn RD et al. Angiotensin II type 1 receptor blockade attenuates TGF-β–induced failure of muscle regeneration in multiple myopathic states. Nat Med 2007;13(2):204–210.

11. Habashi JP et al. Angiotensin II Type 2 Receptor Signaling Attenuates Aortic Aneurysm in Mice Through ERK Antagonism. Science 2011;332(6027):361–365.

12. MacFarlane EG et al. Lineage-specific events underlie aortic root aneurysm pathogenesis in Loeys-Dietz syndrome. J. Clin. Invest. 2019;129(2):659–675.

13. Chen X et al. Conundrum of angiotensin II and TGF-β interactions in aortic aneurysms. Curr Opin Pharmacol 2013;13(2):180–185.

14. Chen X et al. TGF-β Neutralization Enhances AngII-Induced Aortic Rupture and Aneurysm in Both Thoracic and Abdominal Regions. PLoS ONE 2016;11(4):e0153811.

15. Wang Y et al. TGF-beta activity protects against inflammatory aortic aneurysm progression and complications in angiotensin II-infused mice. J. Clin. Invest. 2010;120(2):422–432.

16. Loeys BL et al. Aneurysm syndromes caused by mutations in the TGF-beta receptor. N. Engl. J. Med. 2006;355(8):788–798.

17. Lindsay ME et al. Loss-of-function mutations in TGFB2 cause a syndromic presentation of thoracic aortic aneurysm. Nature Publishing Group 2012;44(8):922–927.

18. Regalado ES et al. Exome sequencing identifies SMAD3 mutations as a cause of familial thoracic aortic aneurysm and dissection with intracranial and other arterial aneurysms. Circ. Res. 2011;109(6):680–686.

19. Tan CK et al. SMAD3 deficiency promotes inflammatory aortic aneurysms in angiotensin II-infused mice via activation of iNOS. J Am Heart Assoc 2013;2(3):e000269.

20. Dai X et al. SMAD3 deficiency promotes vessel wall remodeling, collagen fiber reorganization and leukocyte infiltration in an inflammatory abdominal aortic aneurysm mouse model. Sci Rep 2015;5(1):10180.

21. Doyle AJ et al. Mutations in the TGF-β repressor SKI cause Shprintzen-Goldberg syndrome with aortic aneurysm. Nature Publishing Group 2012;44(11):1249–1254.

22. Carmignac V et al. In-Frame Mutations in Exon 1 of SKI Cause Dominant Shprintzen-Goldberg Syndrome. The American Journal of Human Genetics 2012;91(5):950–957.

23. Akiyoshi S et al. c-Ski acts as a transcriptional co-repressor in transforming growth factor-beta signaling through interaction with smads. J. Biol. Chem. 1999;274(49):35269–35277.

24. Luo K et al. The Ski oncoprotein interacts with the Smad proteins to repress TGFbeta signaling. Genes & Development 1999;13(17):2196–2206.

25. Stroschein SL et al. Negative feedback regulation of TGF-beta signaling by the SnoN oncoprotein. Science 1999;286(5440):771–774.

26. Sun Y et al. Interaction of the Ski oncoprotein with Smad3 regulates TGF-beta signaling. Mol. Cell 1999;4(4):499–509.

27. Xu W et al. Ski acts as a co-repressor with Smad2 and Smad3 to regulate the response to type beta transforming growth factor. Proc. Natl. Acad. Sci. U.S.A. 2000;97(11):5924–5929.

28. Wu JW et al. Structural mechanism of Smad4 recognition by the nuclear oncoprotein Ski: insights on Ski-mediated repression of TGF-beta signaling. Cell 2002;111(3):357–367.

29. Liu X et al. Ski/Sno and TGF-beta signaling. Cytokine Growth Factor Rev. 2001;12(1):1–8.

30. Chen W et al. Competition between Ski and CREB-binding protein for binding to Smad proteins in transforming growth factor-beta signaling. J. Biol. Chem. 2007;282(15):11365– 11376.

31. Gori I et al. Mutations in SKI in Shprintzen-Goldberg syndrome lead to attenuated TGF-β responses through SKI stabilization. Elife 2021;10. doi:10.7554/eLife.63545

32. Holtwick R et al. Smooth muscle-selective deletion of guanylyl cyclase-A prevents the acute but not chronic effects of ANP on blood pressure. Proc. Natl. Acad. Sci. U.S.A. 2002;99(10):7142–7147.

33. Creyghton MP et al. Histone H3K27ac separates active from poised enhancers and predicts developmental state. Proc. Natl. Acad. Sci. U.S.A. 2010;107(50):21931–21936.

34. Rada-Iglesias A et al. A unique chromatin signature uncovers early developmental enhancers in humans. Nature 2011;470(7333):279–283.

35. Visel A et al. ChIP-seq accurately predicts tissue-specific activity of enhancers. Nature 2009;457(7231):854–858.

36. Wang Q et al. Spatial and temporal recruitment of androgen receptor and its coactivators involves chromosomal looping and polymerase tracking. Mol. Cell 2005;19(5):631–642.

37. Jin Q et al. Distinct roles of GCN5/PCAF-mediated H3K9ac and CBP/p300-mediated H3K18/27ac in nuclear receptor transactivation. EMBO J. 2011;30(2):249–262.

38. Weinert BT et al. Time-Resolved Analysis Reveals Rapid Dynamics and Broad Scope of the CBP/p300 Acetylome. Cell 2018;174(1):231–244.e12.

39. Raisner R et al. Enhancer Activity Requires CBP/P300 Bromodomain-Dependent Histone H3K27 Acetylation. CellReports 2018;24(7):1722–1729.

40. Inoue Y et al. Smad3 is acetylated by p300/CBP to regulate its transactivation activity. Oncogene 2007;26(4):500–508.

41. Bowers EM et al. Virtual ligand screening of the p300/CBP histone acetyltransferase: identification of a selective small molecule inhibitor. Chem Biol 2010;17(5):471–482.

42. Kaartinen V, Warburton D. Fibrillin controls TGF-beta activation. Nat Genet 2003;33(3):331–332.

43. Cannaerts E et al. TGF-β signalopathies as a paradigm for translational medicine. Eur J Med Genet 2015;58(12):695–703.

44. Ramirez F et al. Fibrillin microfibrils: multipurpose extracellular networks in organismal physiology. Physiol Genomics 2004;19(2):151–154.

45. Bertoli-Avella AM et al. Mutations in a TGF-β ligand, TGFB3, cause syndromic aortic aneurysms and dissections. J. Am. Coll. Cardiol. 2015;65(13):1324–1336.

46. Courtois A et al. A novel SMAD3 mutation caused multiple aneurysms in a patient without osteoarthritis symptoms. Eur J Med Genet 2017;60(4):228–231.

47. Micha D et al. SMAD2 Mutations Are Associated with Arterial Aneurysms and Dissections. Hum. Mutat. 2015;36(12):1145–1149.

48. Robinson PN et al. Shprintzen-Goldberg syndrome: fourteen new patients and a clinical analysis. Am. J. Med. Genet. 2005;135(3):251–262.

49. Zhang B et al. A dynamic H3K27ac signature identifies VEGFA-stimulated endothelial enhancers and requires EP300 activity. Genome Research 2013;23(6):917–927.

50. Yuan H et al. Involvement of p300/CBP and epigenetic histone acetylation in TGF-β1-mediated gene transcription in mesangial cells. Am. J. Physiol. Renal Physiol. 2013;304(5):F601–13.

51. Petruk S et al. Stepwise histone modifications are mediated by multiple enzymes that rapidly associate with nascent DNA during replication. Nat Commun 2013;4:2841.

52. Dias JD et al. Methylation of RNA polymerase II non-consensus Lysine residues marks early transcription in mammalian cells. Elife 2015;4:387.

53. Huang W-S et al. CIL-102-Induced Cell Cycle Arrest and Apoptosis in Colorectal Cancer Cells via Upregulation of p21 and GADD45. PLoS ONE 2017;12(1):e0168989.

54. Chen G et al. SREBP-1 is a novel mediator of TGFβ1 signaling in mesangial cells. J Mol Cell Biol 2014;6(6):516–530.

55. Santer FR et al. Inhibition of the acetyltransferases p300 and CBP reveals a targetable function for p300 in the survival and invasion pathways of prostate cancer cell lines. Mol. Cancer Ther. 2011;10(9):1644–1655.

56. Marek R et al. Paradoxical enhancement of fear extinction memory and synaptic plasticity by inhibition of the histone acetyltransferase p300. J. Neurosci. 2011;31(20):7486–7491.

57. Maddox SA et al. p300/CBP histone acetyltransferase activity is required for newly acquired and reactivated fear memories in the lateral amygdala. Learn. Mem. 2013;20(2):109–119.

58. Su H et al. Histone Acetyltransferase p300 Inhibitor Improves Coronary Flow Reserve in SIRT3 (Sirtuin 3) Knockout Mice. J Am Heart Assoc 2020;9(18):e017176.

59. Reddy MA et al. Losartan reverses permissive epigenetic changes in renal glomeruli of diabetic db/db mice. Kidney Int. 2014;85(2):362–373.

60. Hayashi K et al. Renin-angiotensin blockade resets podocyte epigenome through Kruppel-like Factor 4 and attenuates proteinuria. Kidney Int. 2015;88(4):745–753.

61. Doyle JJ et al. A deleterious gene-by-environment interaction imposed by calcium channel blockers in Marfan syndrome. Elife 2015;4:e81743.

