## Supplementary Figures for "Epigenetic modulation in the pathogenesis and treatment of inherited aortic aneurysm conditions"

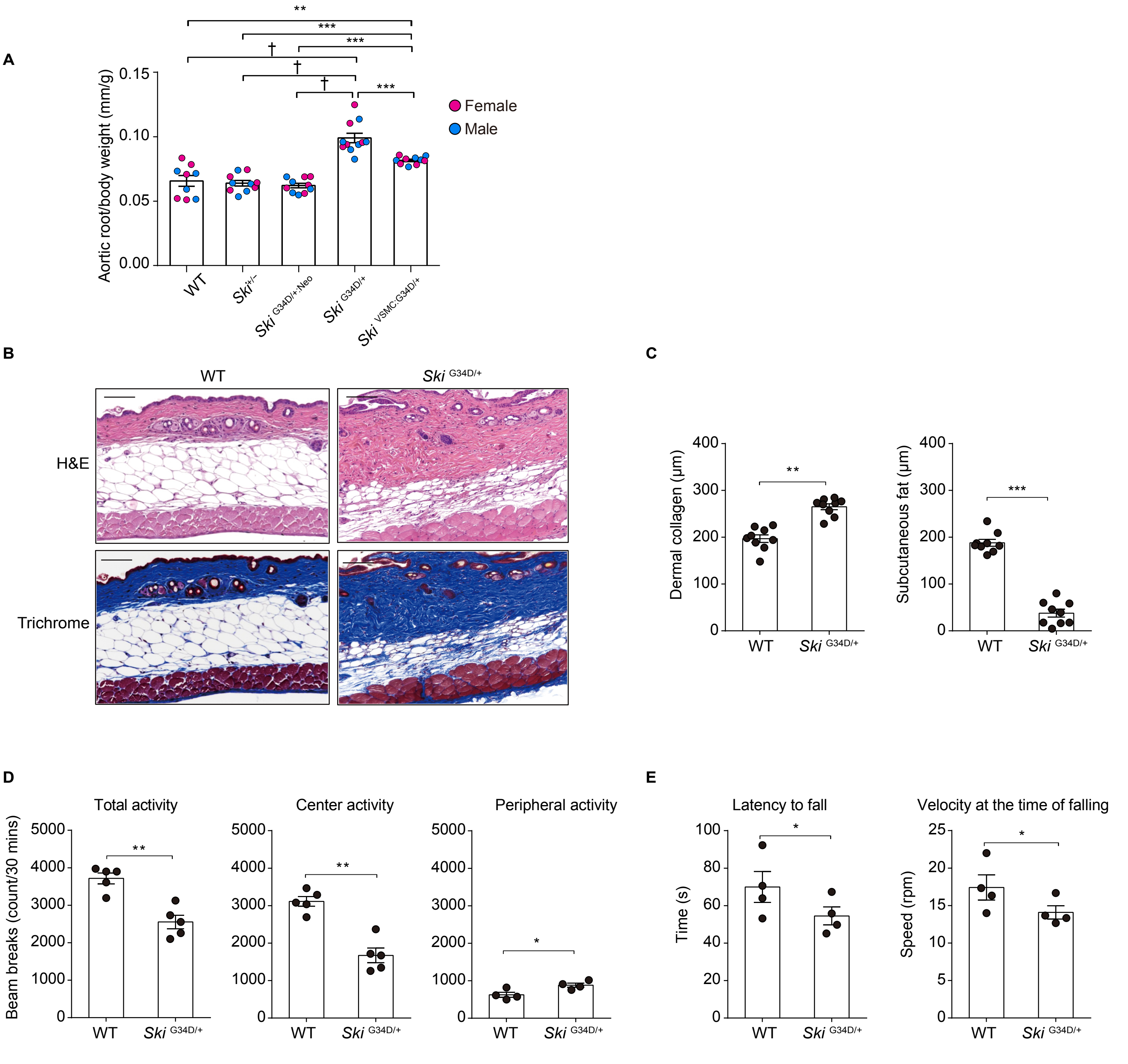


**Figure S1. Aortic, Dermal and neurodevelopmental phenotype of SGS mice.**
(**A**) Mean absolute aortic root diameter normalized to body weight (±SEM) at 24 weeks of age. Note that *Ski*$^{G34D/+}$ and *Ski*$^{VSMC:G34D/+}$ mice had a significantly larger aortic root diameter compared to other genotypes; wild type (WT) (n=9), *Ski*$^{+/-}$ (n=10), *Ski*$^{G34D/+:Neo}$ (n=10), *Ski*$^{G34D/+}$ (n=11), *Ski*$^{VSMC:G34D/+}$ (n=10). (**B**) Dermal phenotype of *Ski*$^{G34D/+}$ mice at 24 weeks of age shown by photomicrographs of representative dorsal dermal sections of *Ski*$^{G34D/+}$ mice and WT littermates stained with hematoxylin and eosin (H&E) and Masson's trichrome stains. (**C**) Dermal collagen thickness and subcutaneous fat layer thickness (±SEM). Compared with WT littermates, *Ski*$^{G34D/+}$ mice demonstrated an increased dermal collagen and a reduced subcutaneous fat layer; *n*=6 in all groups. (**D**) The open-field test results of *Ski*$^{G34D/+}$ mice at 10 weeks old. Compared with WT littermates, *Ski*$^{G34D/+}$ mice exhibited behavioral hypoactivity. The data are shown as beam-break counts per 30 minutes (± SEM). Wild type ($n$ = 5) and *Ski*$^{G34D/+}$ ($n$ = 5). (**E**) The rotarod test results of WT littermates and *Ski*$^{G34D/+}$ mice at 10 weeks old. Compared with WT littermates, *Ski*$^{G34D/+}$ mice demonstrated impaired motor performance. The latency to fall is shown as the time (in seconds ± SEM) that mice remained on an accelerating rotarod (1–20 rpm over 100 seconds) before falling. Non-parametric Kruskal-Wallis test with Dunn's multiple comparison test was used to assess for statistical significance between comparing groups. For all graphs, each bar defines the median with standard error indicated by whiskers and numerical data are presented as scatter dot-plots.*$P$ < 0.05; **$P$ < 0.01; ***P < 0.001; †P < 10-4.

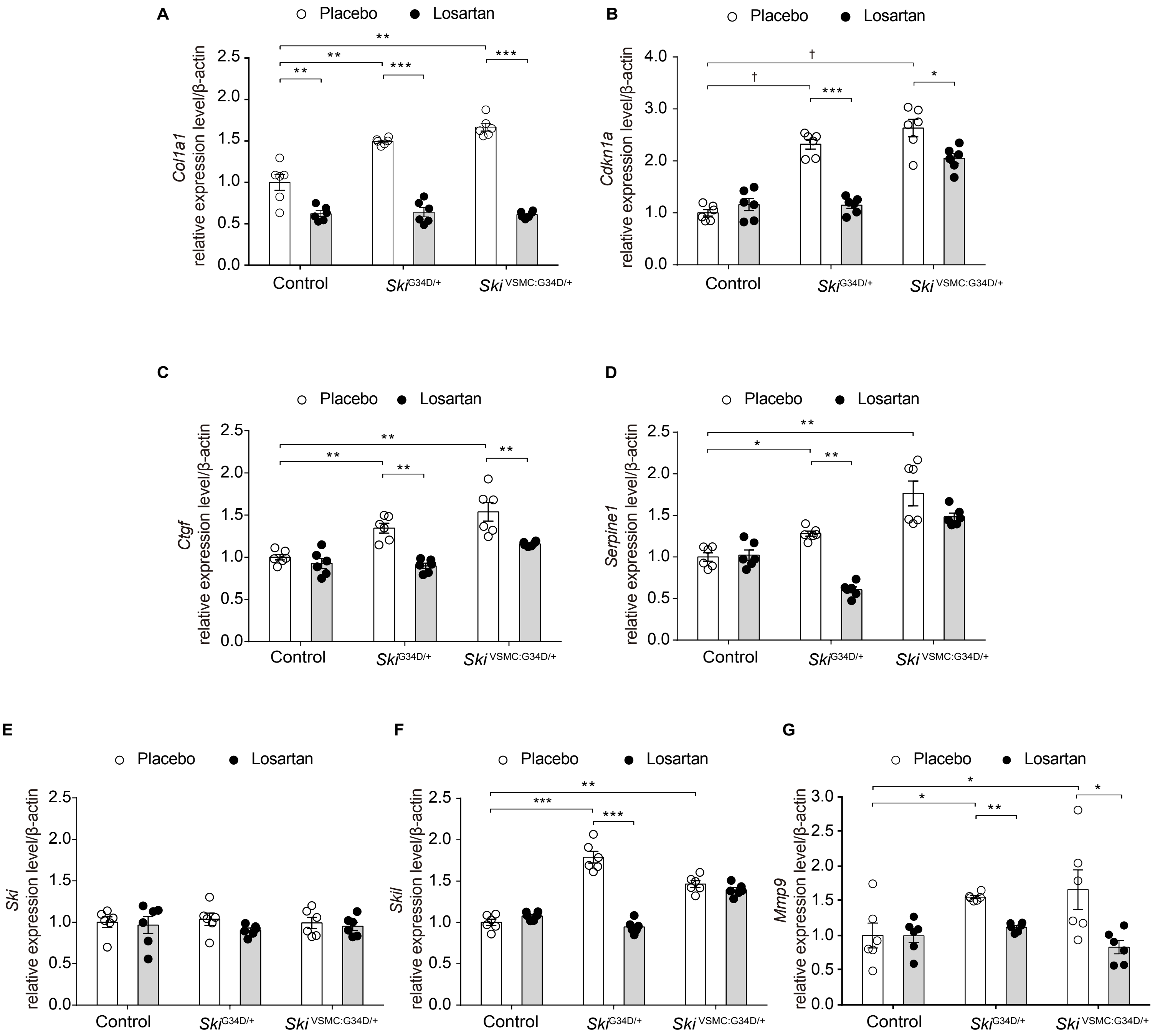


**Figure S2. Expression levels of TGFβ target genes in aortic tissue of SGS mice with Losartan treatment.** Expression levels of TGFβ target genes (A) *Col1a1*, (B) *Cdkn1a*, (C) *Ctgf*, (D) *Serpine1*, (E) *Ski*, (F) *Skil*, and (G) *Mmp9* relative to that of β-actin and normalized by WT expression levels (± SEM), as determined by qPCR. Compared with WT littermates, $Ski^{G34D/+}$ mice and $Ski^{VSMC:G34D/+}$ mice demonstrated increased expression of TGFβ target genes, which was significantly reduced in losartan-treated animals(50 mg/kg/day); n=6 per group. Non-parametric Kruskal-Wallis test with Dunn's multiple comparison test was used to assess for statistical significance between comparing groups. For all graphs, each bar defines the median with standard error indicated by whiskers and numerical data are presented as scatter dot-plots. **P* < 0.05; ***P* < 0.01; ****P* < 0.001; †P < $10^{-4}$.

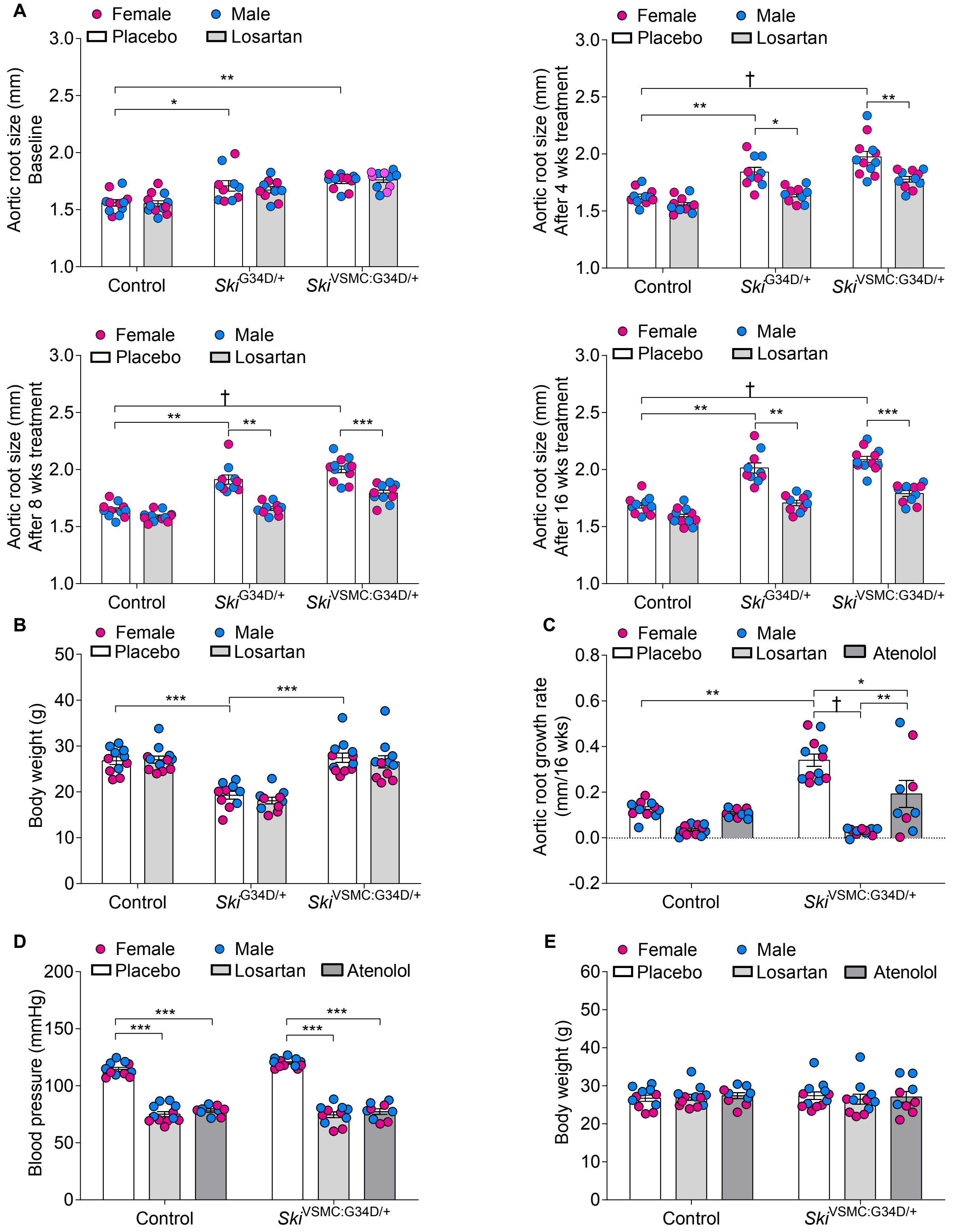


**Figure S3. Aortic root size, body weight and blood pressure of SGS mice after antihypertensive drug treatment.** (**A**) The average absolute aortic root diameter (± SEM) over 16 weeks of treatment with placebo or losartan (50 mg/kg/day) in control *Ski*$^{G34D/+}$ and *Ski*$^{VSMC:G34D/+}$ mice. Control–placebo (*n* = 11), control–losartan (*n* = 12), *Ski*$^{G34D/+}$–placebo (*n* = 10), *Ski*$^{G34D/+}$–losartan (*n* = 10), *Ski*$^{VSMC:G34D/+}$–placebo (*n* = 12), and *Ski*$^{VSMC:G34D/+}$ mice–losartan (*n* = 11). (**B**) Average body weight of control, *Ski*$^{G34D/+}$ and *Ski*$^{VSMC:G34D/+}$ mice after treatment with placebo and losartan for 16 weeks. There was no significant difference in the body weights of placebo- and losartan-treated mice for any genotype. (**C**) Average aortic root growth (± SEM) over 16 weeks of treatment in control and *Ski*$^{VSMC:G34D/+}$ mice with placebo, losartan (50 mg/kg/day), and atenolol (60 mg/kg/day). Control–placebo (*n* = 11), control–losartan (*n* = 12), control–atenolol (*n* = 9), *Ski*$^{VSMC:G34D/+}$–placebo (*n* = 12), *Ski*$^{VSMC:G34D/+}$–losartan (*n* = 11), and *Ski*$^{VSMC:G34D/+}$ mice–atenolol (*n* = 9). (**D**) Average systolic blood pressure in placebo-, losartan-, and atenolol-treated control and *Ski*$^{VSMC:G34D/+}$ mice. There was no significant difference in systolic blood pressure between losartan-treated and atenolol-treated mice. (**E**) Body weights of control and *Ski*$^{VSMC:G34D/+}$ mice after placebo, losartan, and atenolol treatment for 16 weeks. There was no significant difference in the body weights of placebo-, losartan- and atenolol-treated mice for any genotype. Non-parametric Kruskal-Wallis test with Dunn's multiple comparison test was used to assess for statistical significance between comparing groups. For all graphs, each bar defines the median with standard error indicated by whiskers and numerical data are presented as scatter dot-plots. *$P$ < 0.05; **$P$ < 0.01; ***$P$ < 0.001; †$P < 10^{-4}$.

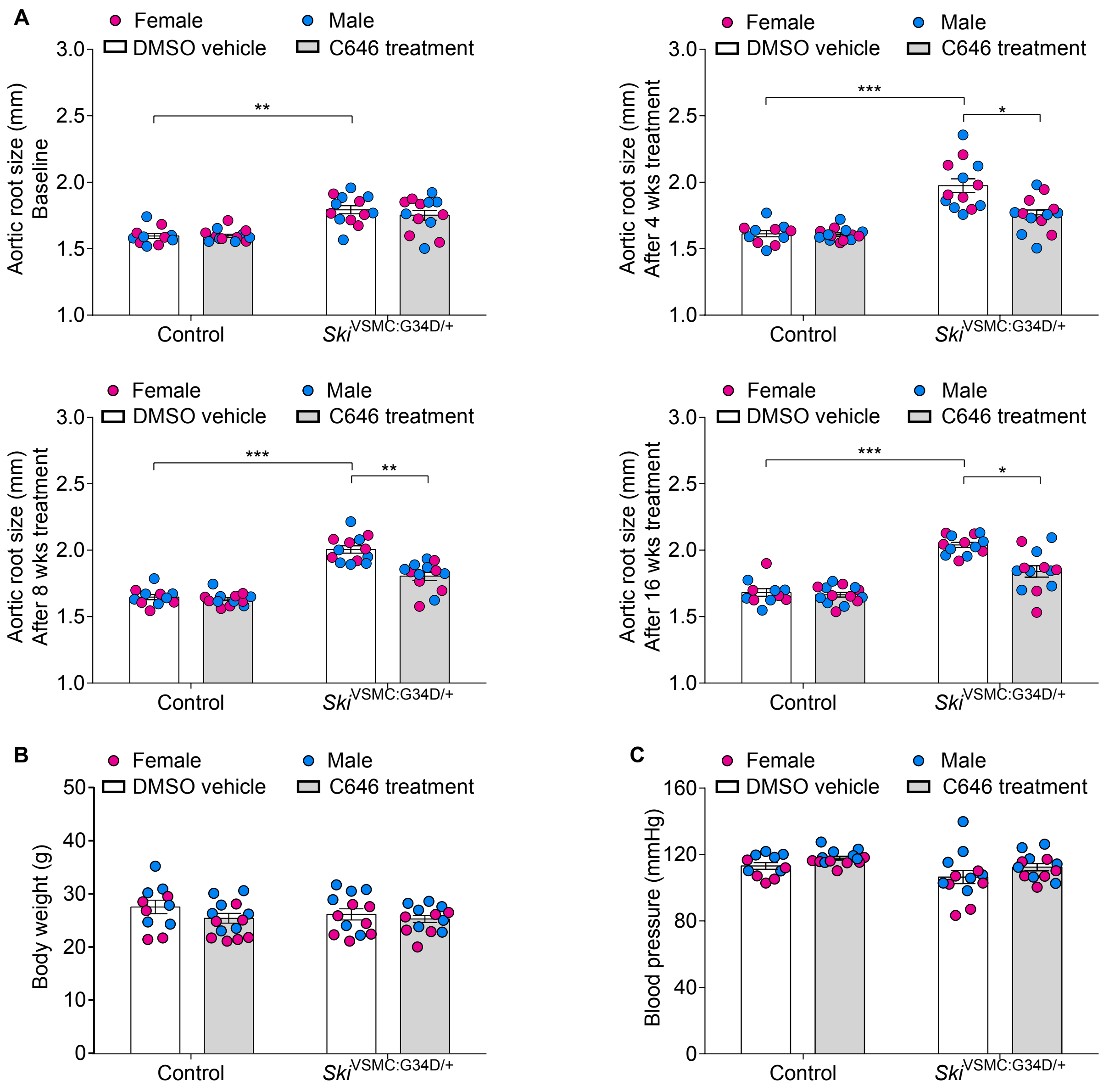


**Figure S4. Aortic root size, body weight and blood pressure of SGS mice after C646 treatment.**
(**A**) Average absolute aortic root diameter (± SEM) over 12 weeks of treatment in control and *Ski*$^{VSMC:G34D/+}$ mice with DMSO (vehicle) and C646 (1mg/kg/day). Control–DMSO (*n* = 11), control–C646 (*n* = 14), *Ski*$^{VSMC:G34D/+}$–DMSO (*n* = 13), and *Ski*$^{VSMC:G34D/+}$–C646 (*n* = 13). (**B**) Body weights of control and *Ski*$^{VSMC:G34D/+}$ mice after placebo and C646 treatment for 12 weeks. There was no significant difference in the body weights of placebo- and C646-treated mice for any genotype. (**C**) Average systolic blood pressure in vehicle DMSO- and C646-treated control and *Ski*$^{VSMC:G34D/+}$ mice. There was no significant difference in systolic blood pressure between vehicle DMSO- and C646-treated mice.Non-parametric Kruskal-Wallis test with Dunn's multiple comparison test was used to assess for statistical significance between comparing groups. For all graphs, each bar defines the median with standard error indicated by whiskers and numerical data are presented as scatter dot-plots. *$P < 0.05$; **$P < 0.01$; ***$P < 0.001$.

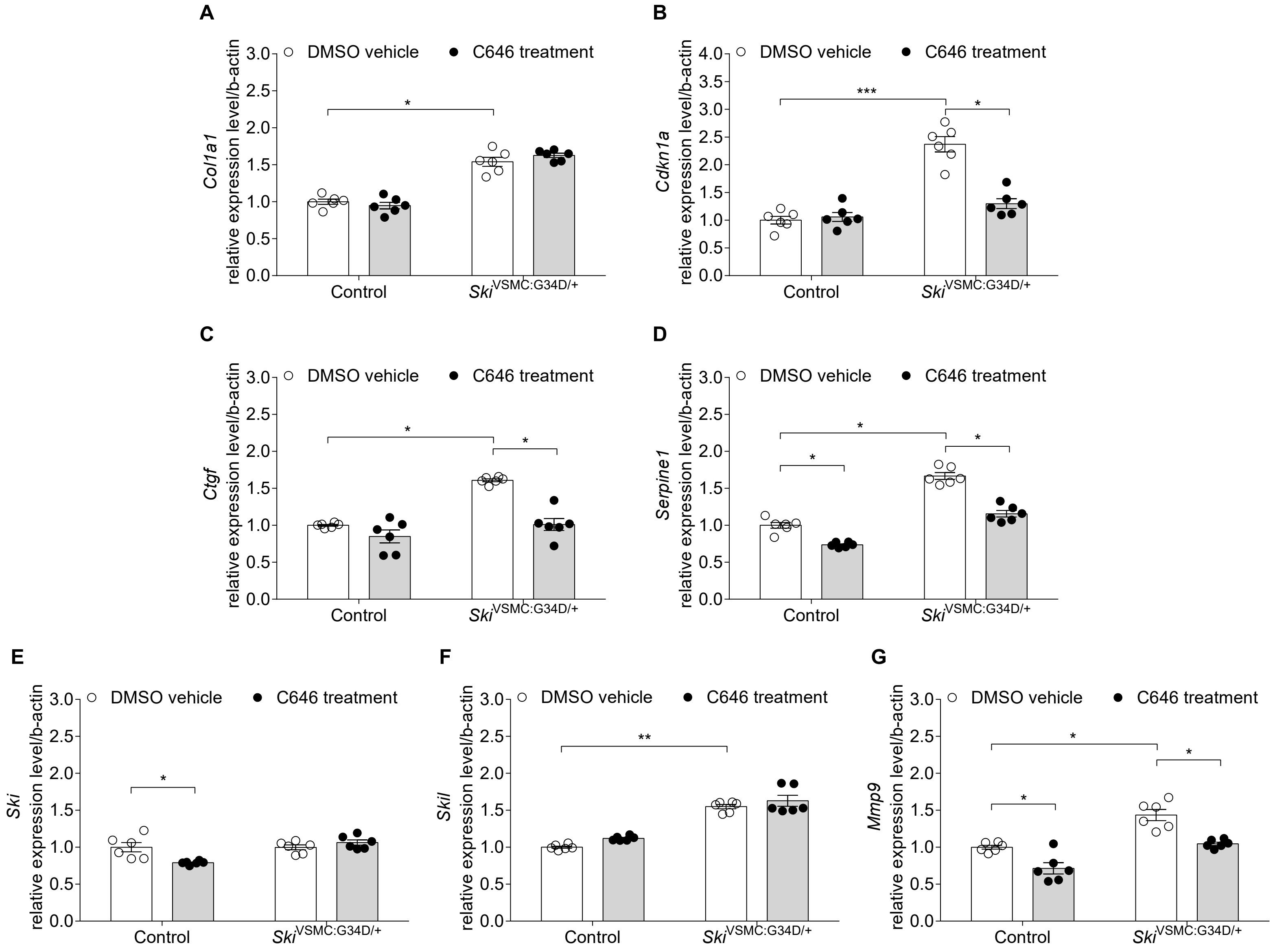


**Figure S5. Expression levels of TGFβ target genes in aortic tissue of SGS mice with C646 treatment.** Expression levels of TGFβ target genes (**A**) *Col1a1*, (**B**) *Cdkn1a*, (**C**) *Ctgf*, (**D**) *Serpine1*, (**E**) *Ski*, (**F**) *Skil,* and (**G**) *Mmp9* relative to that of β-actin and normalized by WT expression levels (± SEM), as determined by qPCR. Compared with control and $Ski^{VSMC:G34D/+}$ mice demonstrated increased expression of TGFβ target genes, which was significantly reduced in C646-treated animals; $n = 6$ per group. $^{*}P < 0.05$; $^{**}P < 0.01$; $^{***}P < 0.001$.

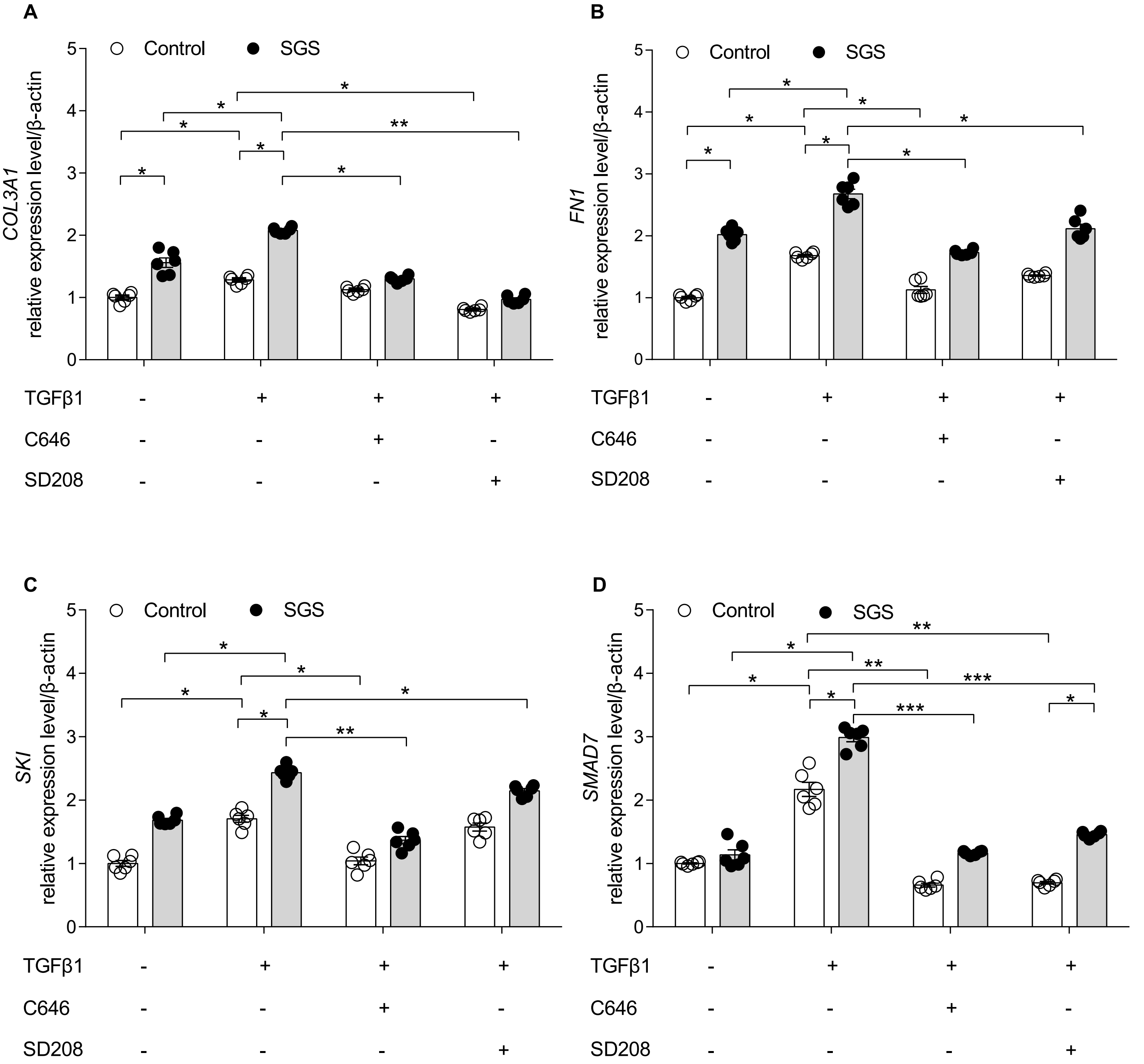


**Figure S6. Expression levels of TGFβ target genes in human fibroblast after C646 treatment.**
TGFβ target gene expression levels in the fibroblasts of controls, and patients with SGS treated with C646 and SD208. Expression levels of (**A**) *COL3A1*, (**B**) *FN1*, (**C**) *SKI*, and (**D**) *SMAD7* relative to that of β-actin in the fibroblasts of patients with SGS, normalized to expression levels of control human fibroblast (± SEM), as determined by qPCR. *$P$ < 0.05; **$P$ < 0.01; ***$P$ < 0.001.
